# A widely-distributed HIV-1 provirus elimination assay to evaluate latency-reversing agents *in vitro*

**DOI:** 10.1101/842419

**Authors:** Kouki Matsuda, Saiful Islam, Toru Takada, Kiyoto Tsuchiya, Benjy Jek Yang Tan, Shin-ichiro Hattori, Hiroo Katsuya, Kosaku Kitagawa, Kwang Su Kim, Misaki Matsuo, Nicole S. Delino, Hiroyuki Gatanaga, Kazuhisa Yoshimura, Shuzo Matsushita, Hiroaki Mitsuya, Shingo Iwami, Yorifumi Satou, Kenji Maeda

## Abstract

Persistence of HIV-1 latent reservoir cells during antiretroviral therapy is a major obstacle for curing HIV-1. Latency-reversing agents (LRAs) are under development to reactivate and eradicate latently infected cells; however, there are few useful models for evaluating LRA activity *in vitro*. Here, we established a long-term cell culture system harboring thousands of different HIV-1-infected cell clones with a wide distribution of HIV-1 provirus similar to that observed *in vivo*. A combination of an LRA and antiretroviral therapy (ART) significantly reduced viral rebound upon treatment interruption. Experimental investigation and mathematical modeling demonstrated that addition of LRA to ART induced latency-reversing effect and contributed to the eradication of replication competent HIV-1. The widely distributed intact provirus elimination (WIPE) assay will be useful for optimizing therapeutics against HIV-1 latency and investigating mechanistic insights into the clonal selection of heterogeneous HIV-1-infected cells.

## INTRODUCTION

Advances in antiviral therapy have dramatically improved the therapeutic options available for treating human immunodeficiency virus type 1 (HIV-1) infection. However, even with the potent combined antiretroviral therapy (cART), HIV-1-infected individuals require lifelong ART because HIV-1 persists in viral reservoirs *in vivo* (*1-3*). The “shock and kill” approach, which first activates cells latently infected with HIV-1 (*4, 5*) using HIV-1 latency-reversing agents (LRAs), induces viral production, and eliminates virus-producing cells by immune clearance or viral cytopathic effects, is a possible strategy for curing HIV-1 (*6-8*). Previous studies demonstrated that LRAs were potent in *in vitro* assays but were not necessarily effective *in vivo* because characteristic of the viral reservoir is quite different between *in vitro* and *in vivo (9)*. HIV integration site significantly impact the proliferation of HIV-infected cells. Yet, no *in vitro* system recapitulate the heterogeneity of HIV-infected cells *in vivo* (*10-15*).

Currently, there are three major model systems which can provide preclinical testing of HIV cure strategies: *in vitro* primary cell or cell line systems, *ex vivo* testing in clinical samples, and *in vivo* animal models. The key determinants of the success of these models is whether they can recapitulate drug effects *in vivo*. In particular, genetic and epigenetic environments are key factors that determine the fate of HIV-1 provirus, either active viral production or viral latency (*13*). Currently available *in vitro* models for HIV-1 latency, such as ACH2, J1.1, and U1 cells, carry just only one or two proviruses integrated into particular host genomic regions, and cannot recapitulate the thousands of different integration sites observed *in vivo* (*10, 11*). Therefore, latency reversing agents (LRAs) tested in *in vitro* model systems fail to demonstrate responses *ex vivo* (*9*) or *in vivo* (*16*). Primary cell models recapitulate the diverse integration sites (*17-19*), but most cells die during lengthy cultures (*20*). *Ex vivo* models using clinical samples from HIV-1-infected individuals may also capitulate the diverse HIV-1 integration sites (*7*). However, the rarity of HIV-1-infected cells in HIV-1-infected individuals (*14, 21*) prevents scalable and robust analysis of the impact of LRA in respect to HIV-1 integration sites. Although animal models provide valuable pre-clinical testing, the cost and resources required prevents large-scale screening (*22-24*).

In the present study, we aimed to establish a new *in vitro* infection model that harbors a much wider variety of HIV-1-infected clones than that of conventional *in vitro* models. Our *in vitro* model, the widely distributed intact provirus elimination (WIPE) assay, recapitulates the diverse HIV-1 integration sites observed *in vivo*. This new cell-line based *in vitro* system provides not only a robust and scalable system to examine HIV-1 latency reversal but also a new platform to study mechanisms of HIV-1 persistence in respect to the diverse HIV-1 integration sites and the host genomic environment.

## RESULTS

### Development of an *in vitro* model mimicking the distribution of HIV-1 provirus *in vivo*

To establish an HIV-1 chronically-infected *in vitro* model with a wide variety of HIV-1-infected clones, several host cell lines were infected with an HIV-1 infectious clone, HIV-1_NL4-3_ or HIV-1_JRFL_. Cell growth and viral production in the supernatant was monitored twice a week. MT-4 cells infected with HIV-1_NL4-3_ rapidly underwent apoptosis due to vigorous viral production in MT-4 cells, resulting in no live cells remained after 30 days (data not shown). We analyzed intracellular p24 expression in samples (with cell viability of 60-70 %) on day 30. Jurkat cells infected with HIV-1_NL4-3_ (Jurkat/NL) maintained high levels of viral production (Fig. 1A) and cell-associated HIV-1 DNA (Fig. 1B). Hut78 cells and MOLT4 cells infected with HIV-1_NL4-3_ also continued to produce HIV-1, however, the viral production and cell-associated HIV-1 DNA decreased on day 30 (Figs. 1A and 1B). PM1CCR5 cells infected with HIV-1_JRFL_ (PM1CCR5/JRFL) maintained HIV-1 production and intracellular HIV-1 DNA levels after 30 days of culture; however, the p24 level in the supernatant decreased after 90 days (data not shown). Flow cytometry analyses on day 30 revealed four distinct cell populations, including p24-negative live cells, p24-negative dead cells, p24-positive live cells, and p24-positive dead cells (Fig. 1C), in the three tested infected cell lines. Based on these results, we focused on Jurkat/NL in the present study, because that maintained heterogenous HIV-1 infectivity for a long period of time. We confirmed the infectivity of viruses produced in the supernatant of Jurkat/NL cells using MT-4 cells (fig. S1A).

**Fig. 1.**
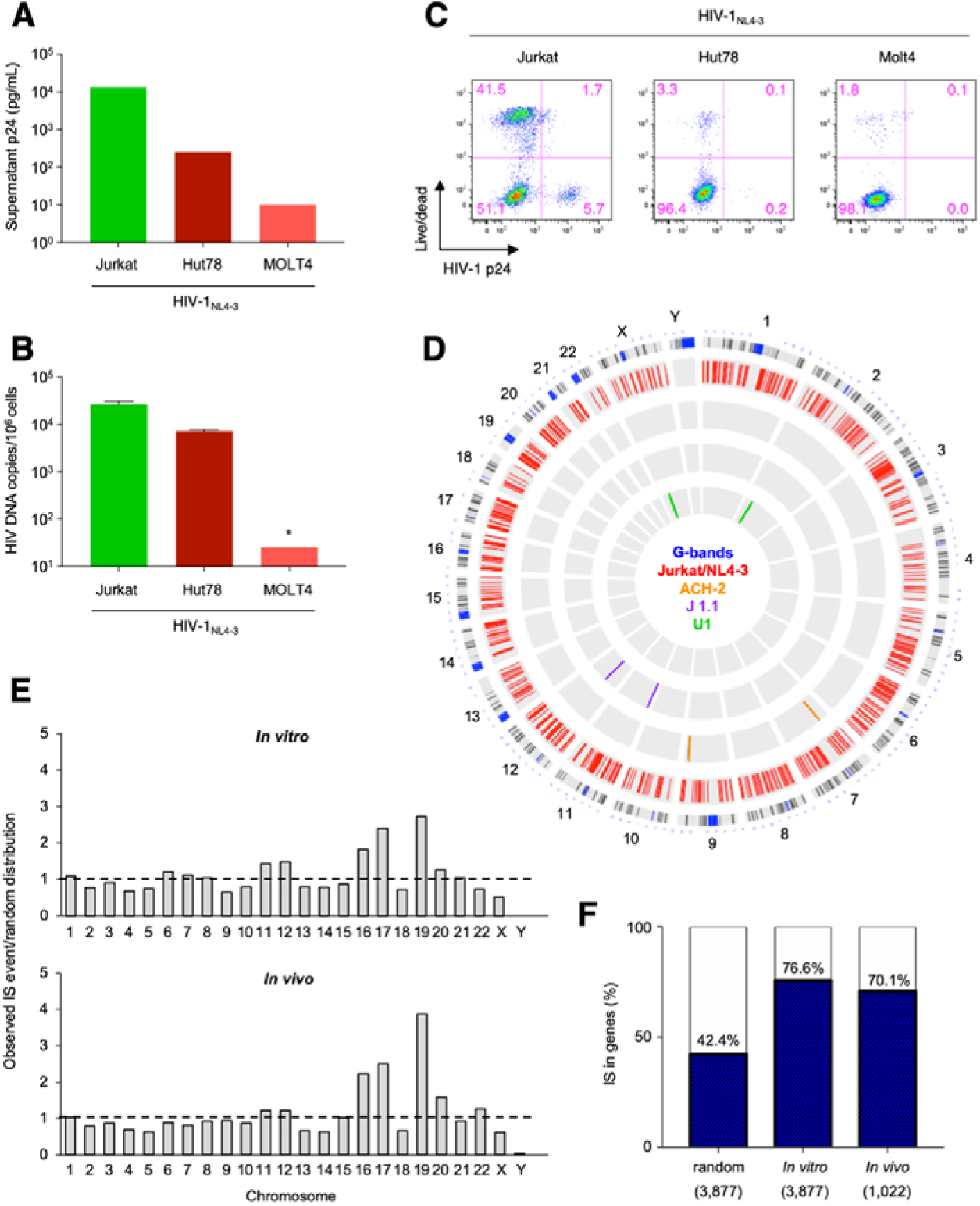
Establishment of a new *in vitro* HIV-1 infection model. To establish a cell culture model of long-term persistent HIV-1 infection, various T-cell lines (Jurkat, Hut78, and MOLT4 cells) were used. HIV-1 production (**A**) and copies of intracellular HIV-1 DNA (**B**) on day 30 of each cell line (**C**) Co-existence of p24-positive and p24-negative cell populations in HIV-1–infected Jurkat, Hut78, and MOLT4 cell lines. The percentage of intracellular p24-positive cells was analyzed by flow cytometry. (**D**) Circos plot depicting viral integration sites (IS) across the human genome in the Jurkat/NL system and in different cell lines *in vitro*. Each chromosome is presented on the outer circle and is broken down into sequential bins. Blue/black, red, orange, purple, and green bars indicate G-bands (condensed chromosome region by Giemsa staining for karyotyping), Jurkat/NL system, ACH-2, J1.1, and U1, respectively. (**E**) Comparison of HIV-1 IS frequency in the individual chromosomes in the *in vitro* model (Jurkat/NL) and *in vivo* in PBMCs from five HIV-1-infected individuals. The *y-*axis depicts the proportion of integration events observed relative to random distribution, with a horizontal dashed black line set at a value of 1. (**F**) Relationship between HIV-1 IS and the host genes, *in vitro* and *in vivo*, compared via random distribution. Numbers in parentheses at the bottom of the bars indicate the numbers of unique ISs observed; numbers at the top of the bars indicate the percentage of HIV-1 proviruses integrated within the host genes in each group. Asterisk (*) stands for below detection limit.

We next analyzed the number of HIV-1-infected clones in the *in vitro* culture model. In principle, each HIV-1-infected clone has a different viral integration site (IS), which can be used to distinguish clones. To determine HIV-1 IS, we performed ligation-mediated polymerase chain reaction (LM-PCR) to detect the junctions between the 3′-long terminal repeat (LTR) of HIV-1 and the flanking host genome sequence (*25, 26*). We analyzed genomic DNAs extracted from an aliquot of Jurkat/NL cells in the culture and detected approximately 2,933 different ISs, demonstrating the presence of thousands of different infected clones in the total culture well (Fig. 1D). This was in stark contrast with ACH-2, J1.1, and U1 cell lines, in which only two ISs were detected (Fig. 1D). Next, we tested whether the distribution of HIV-1 proviruses in the *in vitro* system was comparable to that found *in vivo*. We analyzed HIV-1 ISs in peripheral blood mononuclear cells (PBMCs) isolated from HIV-1-infected individuals following the same protocol as that for *in vitro* cultured cells. We observed several similarities between HIV-1 integration *in vitro* and *in vivo*, i.e., increased integration incidence in certain chromosomes (Fig. 1E). First, we found enrichment of integration in chromatin 16, 17, and 19, similar to that observed in vivo (*27*) (Fig. 1F). Second, we found an enrichment of integration into the host gene bodies (75.7%), similar to those observed *in vivo* (70.8%). (*14, 27, 28*). Third, we found that 48.7 % of integration are in the same orientation as the host transcription unit, similar to that found in vivo (*27, 28*). These data indicate that the new *in vitro* model is far more heterogenous than conventional latent cell line models for HIV-1 and recapitulates the HIV-1 integration sites found *in vivo*.

### The WIPE assay recapitulates the effect of LRA and ART in HIV-infected cells

We next evaluated the effect of antiretroviral agents and/or LRAs against various HIV-1-infected clones in the *in vitro* culture system. First, we evaluated 11 LRAs to determine their activity in Jurkat/NL cells, and found that PEP005 [ingenol-3-angelate, protein kinase C (PKC) activator] induced HIV-1 production and apoptosis in HIV-1-infected cells at the lowest concentration tested (fig. S1B and C). Furthermore, 1 μM SAHA and panobinostat induced strong caspase-3 activation (fig. S1C); however, the same concentration of these drugs also induced strong cell toxicity in HIV-1-negative cells (data not shown). Hence, we used PEP005 in subsequent experiments. We cultured Jurkat/NL cells in the presence or absence of an antiretroviral drug and/or an LRA (Fig. 2A). The assay started with a total of 5 × 10^4^ Jurkat-NL cells in 2 mL culture wells with/without a drug, and the cell number reached around 1 million in wells on day 7. The Jurkat-NL cells contains approximately 5 × 10^4^ copies of HIV-1 DNA in 1 × 10^6^ cells (data not shown). The medium in each well was exchanged every 7 days with the same number (5 × 10^4^) of cells with/without a drug. We used the antiretroviral drug EFdA (4′-ethynyl-2-fluoro-2′-deoxyadenosine)/MK-8591/islatravir (ISL), which is a potent nucleoside reverse-transcriptase inhibitor and currently under clinical trials (*29*). Treatments with EFdA (50 nM), PEP005 (5 nM), or a combination of EFdA and PEP005 were started simultaneously, and supernatant p24 levels were monitored for 4 months (Fig. 2A). Time-course data from multiple independent experiments are shown in Fig. 2B. Treatment with PEP005 alone did not suppress HIV-1 replication during the first 9 weeks of cultivation, while EFdA alone successfully decreased viral production in the supernatant to undetectable levels after 4–6 weeks. The combination treatment with EFdA and PEP005 also decreased the amount of HIV-1 in the supernatant to undetectable levels.

**Fig. 2.**
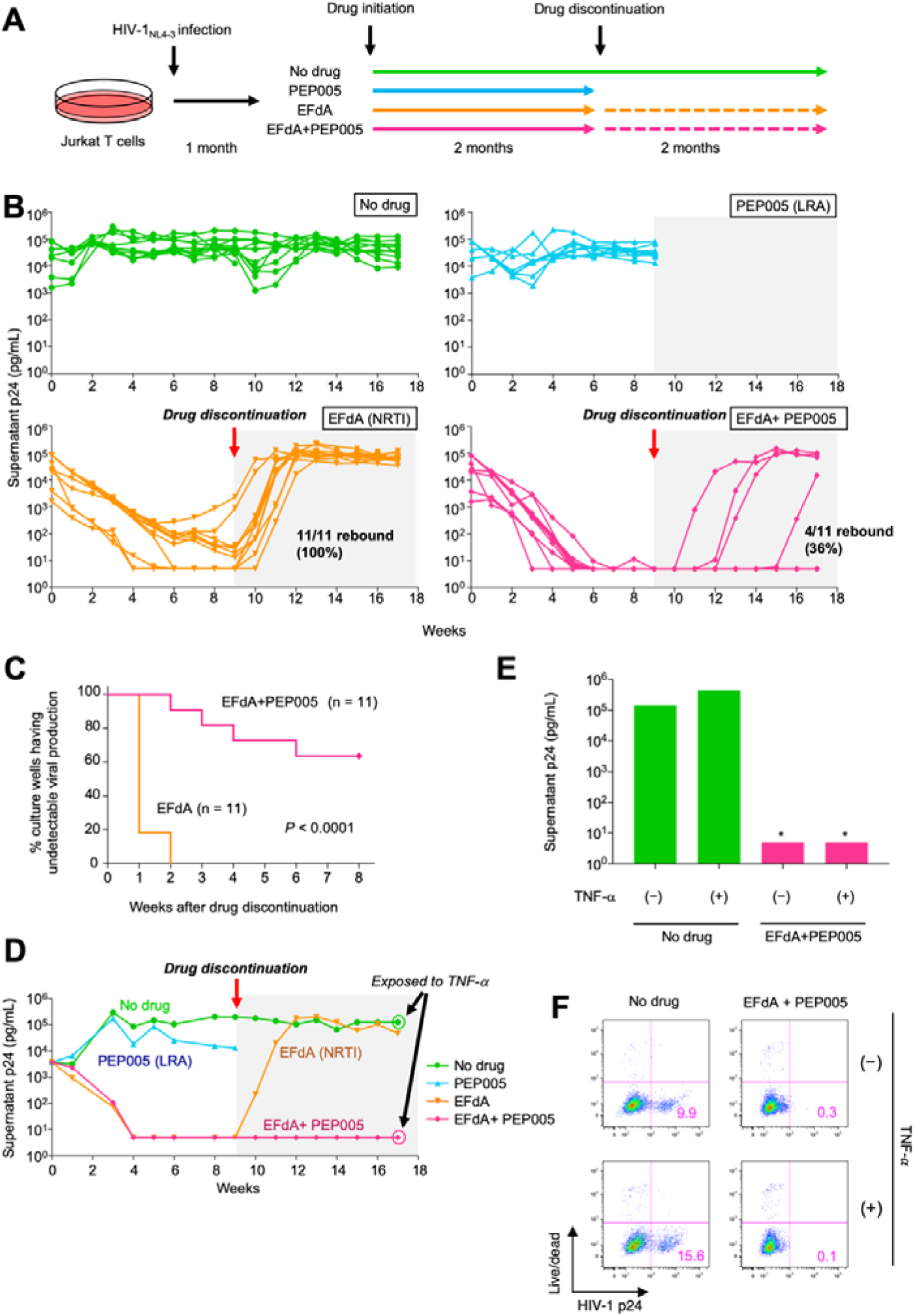
Effect of drug treatments on viral persistence in the new *in vitro* infection model. **(A)** Assay overview. Schematic representation of the assay protocol involving the HIV-1_NL4-3_–infected cell culture model (Jurkat/NL cells). **(B)** Changes in supernatant p24 levels without drugs, with 5 nM PEP005 or 50 nM EFdA, or with a combination of 50 nM EFdA and 5 nM PEP005 (n = 11, 9, 11, and 11, respectively). Drug treatment was terminated on week 9 but analysis continued for an additional 8 weeks. **(C)** Log-rank test comparison of the percentage of non-recurrence in the EFdA single treatment and the combination treatment. **(D)** Changes in supernatant p24 levels in a representative experiment (Exp. 1) from experiments shown in Fig. 2b. **(E–F)** Assessment of the viral rebound in Jurkat/NL cells after drug discontinuation. Cells treated with drugs or untreated cells were stimulated with TNF-α (10 ng/mL) in week 17, and supernatant p24 (**E**) and intracellular p24 levels (**F**) were analyzed on day 6 after stimulation. Asterisk (*) denotes below detection limit.

We next interrupted the drug treatment in week 9 and observed a remarkable rebound of viral production in samples treated with EFdA alone (11/11, 100%); however, no rebound was found in 64% (7/11) of samples treated with the combination of EFdA and PEP005 (Fig. 2B) and the difference was statistically significant (Log-rank test, p < 0.0001) (Fig. 2C). In one representative experiment (Exp. 1), viral rebound was observed in cells treated with EFdA only, but not in cells treated with both EFdA and PEP005 (Fig. 2D). Viral rebound from EFdA + PEP005-treated cells was not detected (in the supernatant or intracellularly; for a total of ∼1 million cells in 2 mL culture wells) even after stimulation with tumor necrosis factor-α (TNF-α) in week 17, confirming lack of replication-competent HIV-1 in the treated sample (Fig. 2E and F). Combinations with other antiretroviral agents, i.e., darunavir (DRV, a protease inhibitor) and dolutegravir (DTG, an integrase inhibitor), or other LRAs (SAHA, an HDAC inhibitor; prostratin, a PKC activator) were also tested (fig. S2). Antiretroviral drugs (DRV, DTG, and EFdA) effectively decreased supernatant HIV-1 levels, whereas most LRAs failed to suppress viral replication when used on their own. However, the combination of an LRA with an antiretroviral drug delayed or inhibited viral rebound after treatment interruption similar to the recent report regarding LRA activity in an *in vivo* model (*30*) and in clinical trials (*31*). Among several drug combinations analyzed in the present study, only the combination of EFdA and PEP005 resulted in a significant reduction in viral rebound. Overall, we showed that WIPE assay is a robust and scalable assay which examines the effect of both ART and LRAs.

### Evidence of reactivation of latently infected cells and cell apoptosis induced by combination treatment containing LRA

The rationale behind using LRA for an HIV-1 cure is reactivating the latent HIV-1 provirus and inducing cell apoptosis via cytopathic effects or recognition by the host antiviral immunity (*7, 32*). We therefore analyzed whether the reduction in viral rebound observed in the present study was indeed mediated by the elimination of latently infected cells. Because Jurkat cells are always proliferating and active, it was generally thought that HIV-infected Jurkat cells will not undergo latency. First, we investigated the presence of latent clones in the Jurkat/NL system by infecting Jurkat-LTR-green fluorescent protein (GFP) cells (Jurkat cells stably transfected with a plasmid containing the GFP reporter gene driven by the HIV-1 promoter LTR) with HIV-1_NL4-3_, in which GFP expression is dependent on the presence of exogenous viral protein Tat. We sorted and stimulated the GFP-negative cell fraction with TNF-α and found that the treatment increased proviral transcription in this fraction (Fig. 3A and B), indicating the presence of latently infected cells in the Jurkat/NL system. The percentage of such latently infected cells was approximately 1% in this assay (Fig. 3C). Since antiviral cytotoxic T-lymphocytes (CTLs) and antibodies are absent in the Jurkat/NL system, latent HIV-1-infected cells reactivated by PEP005 would have been eliminated mainly by viral cytopathicity or cell apoptosis (fig. S1C) (*32, 33*). To analyze relationship between viral protein production and cell apoptosis, we examined intracellular HIV-1 p24 and cell apoptosis by flowcytometry during the early phase of drug treatment. We then found an increase in p24 protein expression in cells treated with PEP005 (+/− EFdA) just 6 h after drug treatment initiation (Fig. 3D), which was followed by an increase in annexin V expression (Fig. 3E). The number of intracellular p24^+^ cells decreased in EFdA- or EFdA + PEP005-treated cell populations, and these cells constituted less than 2.5% of the total population by week 2 (Fig. 3F). On week 4, we analyzed caspase-3 levels in these cells and found that caspase-3 expression was much higher in p24^+^ cells, especially in the EFdA + PEP005-treated cells, than in p24^−^ cells (Fig. 3G). These observations suggest that PEP005 functions as an LRA, inducing apoptosis in cells latently infected with HIV-1 and facilitating the elimination of HIV-1 replicable cells *in vitro*. The proportion of the full-length-type provirus among total proviruses in EFdA alone or EFdA + PEP005-treated cells was also decreased but was nonetheless more than 50% after the initial 3-week treatment (Fig. 3H), suggesting that more than half of all proviruses were thought to be replication-competent at this time point. Collectively, these data indicate that the elimination of HIV-1 producible cells achieved in this study was at least partially due to the reactivation of latent HIV-1 reservoirs.

**Fig. 3.**
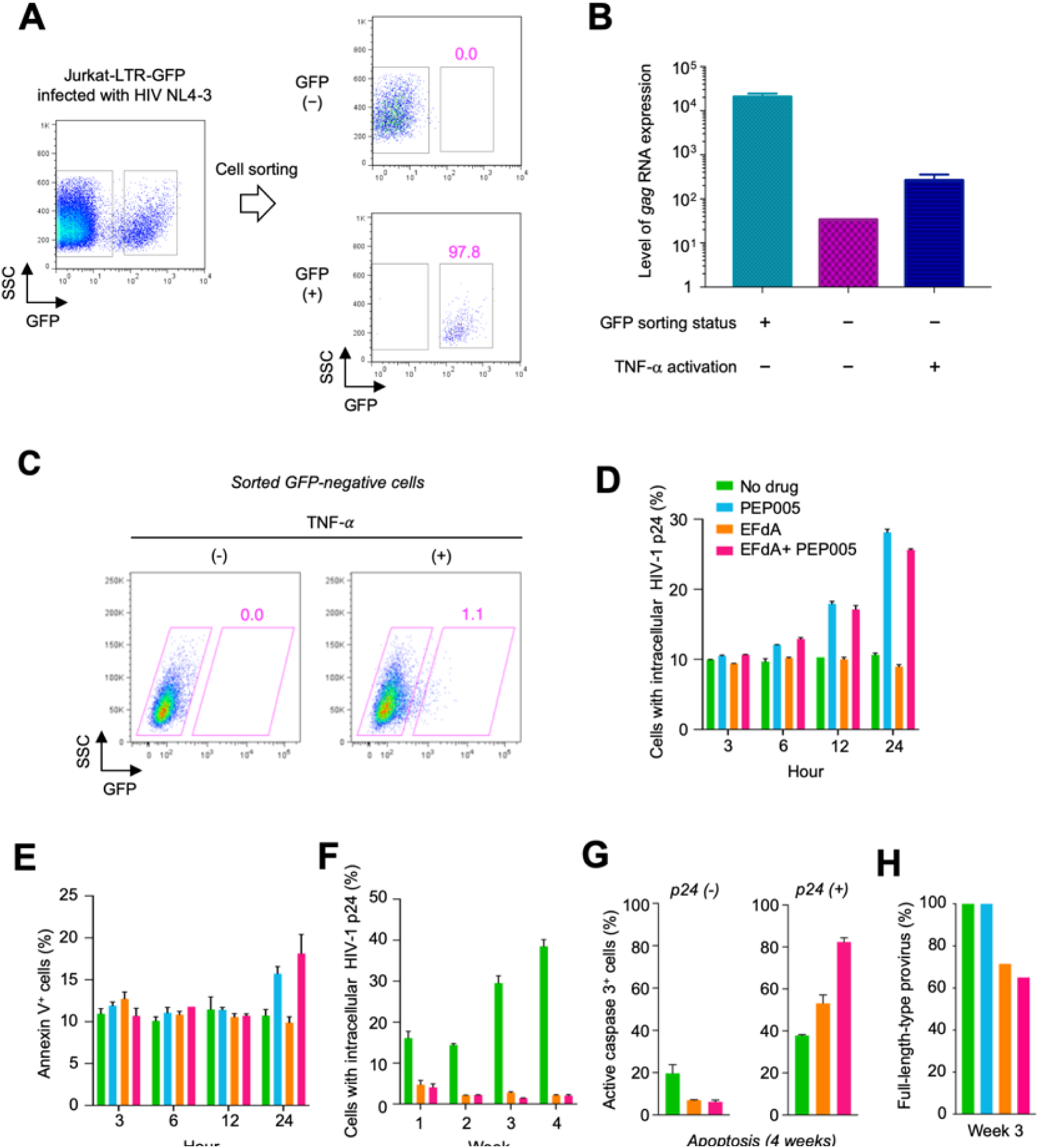
Proof-of-concept of the shock and kill strategy of the WIPE assay. **(A)** Sorting of GFP-positive cells among HIV-1-infected Jurkat-LTR-GFP cells. GFP-positive (Tat^+^) or GFP-negative populations (Tat^-^) among HIV-1-infected Jurkat-LTR-GFP cells were sorted. (**B)** GFP-negative cells were stimulated with 10 ng/mL TNF-α for 6 h and *gag* mRNA expression was quantified. **(C)** GFP expression in sorted GFP-negative Jurkat-LTR-GFP cells infected with HIV-1_NL4-3_ was analyzed after 6 h of TNF-α stimulation (10 ng/mL). **(D**–**E)** p24 expression and cell apoptosis during the early phase of drug treatment. Bar graphs show the change in the percentage of cells expressing intracellular HIV-1 p24 (**D**) and annexin V (**E**) during the initial 24 h of drug treatment. **(F)** Changes in the numbers of cells with intracellular p24 (weeks 1–4). **(G)** Percentages of active caspase-3-positive cells in the p24-positive or p24-negative cell population. **(H)** Percentages of full-length-type HIV-1 provirus after 3 weeks of drug treatment. Data represent the mean ± S.D. of three independent experiments.

### Quantifying reactivation dynamics of latent HIV-1 reservoirs

To analyze how addition of LRA affect the kinetics of HIV-1 replication in the WIPE assay, we developed a mathematical model describing HIV-1 infection dynamics with antiviral drugs (see Supplementary Text). Then, to assess the variability of kinetic parameters and model predictions, we performed Bayesian estimation for the whole dataset using Markov chain Monte Carlo (MCMC) sampling (see Supplementary Text). The typical behavior of the model using these best-fit parameter estimates is shown together with the data in Fig. 4A, and indicated that the mathematical model described the WIPE assay data very well. The shadowed regions corresponded to 95% posterior predictive intervals, the solid lines gave the best-fit solution (mean) for the mathematical model, and the colored dots showed the experimental datasets. Note that our mathematical model does not account defective proviral DNA but our experimental measurement included the defective provirus. Since most of proviral DNA were defective in EFdA + PEP005-treated cell (see Fig.5B and C in next section), our mathematical model which predicts only intact proviral DNA underestimated the normalized proviral load. In fact, our mathematical model well predict the proviral load in EFdA-treated cell since most of proviral DNA were intact.

**Fig. 4.**
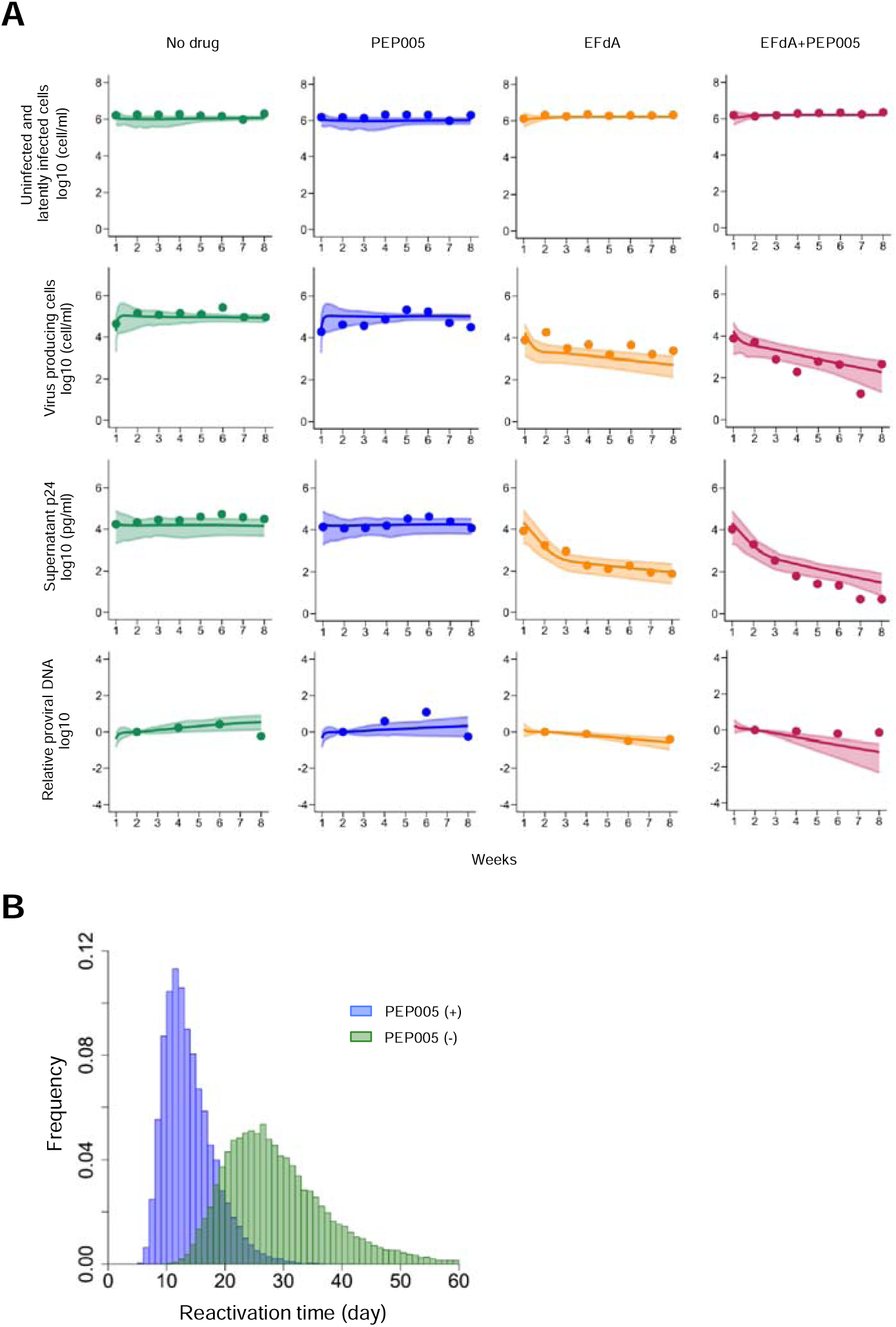
HIV-1 infection dynamics with antiviral drugs in WIPE assay. **(A)** Fitting of the mathematical model to the experimental data in WIPE assay without and with antiviral drug(s): numbers of uninfected and latently infected cells (cells/ml), virus producing cells (cells/ml), supernatant p24 (pg/ml), and normalized proviral DNA. The shadowed regions corresponded to 95% posterior predictive intervals, the solid lines gave the best-fit solution (mean) for the mathematical model, and the colored dots showed the experimental datasets. All data were fitted simultaneously. **(B)** The distribution for the time until reactivation without and with PEP005-treatment calculated from all accepted MCMC parameter estimates are shown in green and blue, respectively. These lengths were significantly shorter with PEP005-treatment than without treatment as assessed by the repeated bootstrap *t*-test.

To evaluate whether PEP005-treatment induced the reactivation of latent HIV-1 reservoirs in WIPE assay, we applied our mathematical model to time-course experimental data. With estimated distributions of parameters *a* and *ε*, we calculated the distribution for the time until reactivation without and with PEP005-treatment (1/*a* and 1/*εa*, respectively) (Fig. 4B). Comparing these lengths without and with PEP005-treatment showed a significant difference (29.1 days, 95% CI: 15.9 − 52.1 days and 13.9 days, 95% CI: 7.85 − 24.3 days, respectively) (*p* = 2.59 × 10^−4^ by repeated bootstrap *t*-test). These estimates indicated that PEP005-treatment induced the reactivation in WIPE assay and significantly reduce the time until reactivation (that is, 52.2% reduction on average).

### Long-term LRA+ART preferentially selects infected clones with defective HIV-1 proviruses

To elucidate the possible mechanism(s) underlying the reduction in viral rebound *in vitro*, we quantitatively and qualitatively analyzed HIV-1 proviruses during time-course of treatment. We analyzed cell-associated HIV-1 DNA loads in one representative experiment (Exp. 6) (fig. S3A) and found that the HIV-1 DNA load was markedly decreased in samples treated with EFdA alone (Fig. 5A). The addition of PEP005 to EFdA further decreased the HIV-1 DNA load (Fig. 5A). After drug discontinuation, the HIV-1 DNA load increased in the sample treated with EFdA alone but not in the sample treated with both drugs (Fig. 5A). We characterized the structure of the HIV-1 proviral genome by nearly full-length PCR, using a single copy of the HIV-1 genome as a template (*34*). We observed an increased proportion of a defective HIV-1 genome after EFdA treatment at 9 weeks, possibly due to preferential elimination of intact, replication-competent proviruses (Fig. 5B and C). But, after drug discontinuation of EFdA treatment at 17 weeks, intact HIV-1 proviruses emerged again, suggesting new rounds of infection from latently infected cells harboring replication competent HIV-1 (Fig. 5B and C). Strikingly, combination of ART (EFdA) and LRA (PEP005) eliminated intact proviruses (Fig. 5B and C), suggesting that LRA has reactivated all cells harboring replication-competent HIV and exposes them for viral cytopathic effect and cell death. All proviruses detected in the sample after a combined EFdA and PEP005 treatment were defective ones at 17 weeks after initiation of drug treatment (Fig. 5B and C). Accumulation of defective proviruses during anti-retroviral therapy has been reported (*14, 34*) and we also confirmed that in this study. We analyzed peripheral blood samples of HIV-1-carrying individuals (table S1) with nearly full-length, single-genome PCR. Before the initiation of cART, 25–42% proviruses in PBMCs from HIV-1 infected individuals were defective, and the ratio increased to 83–100% after successful cART (treatment duration of at least 6 years) (Fig. 5D and fig. S4 A and B). The data suggest that our WIPE assay recapitulated the accumulation of defective proviruses *in vivo* caused by a preferential selection of defective and/or replication-incompetent proviruses during long-term antiretroviral treatment.

**Fig. 5.**
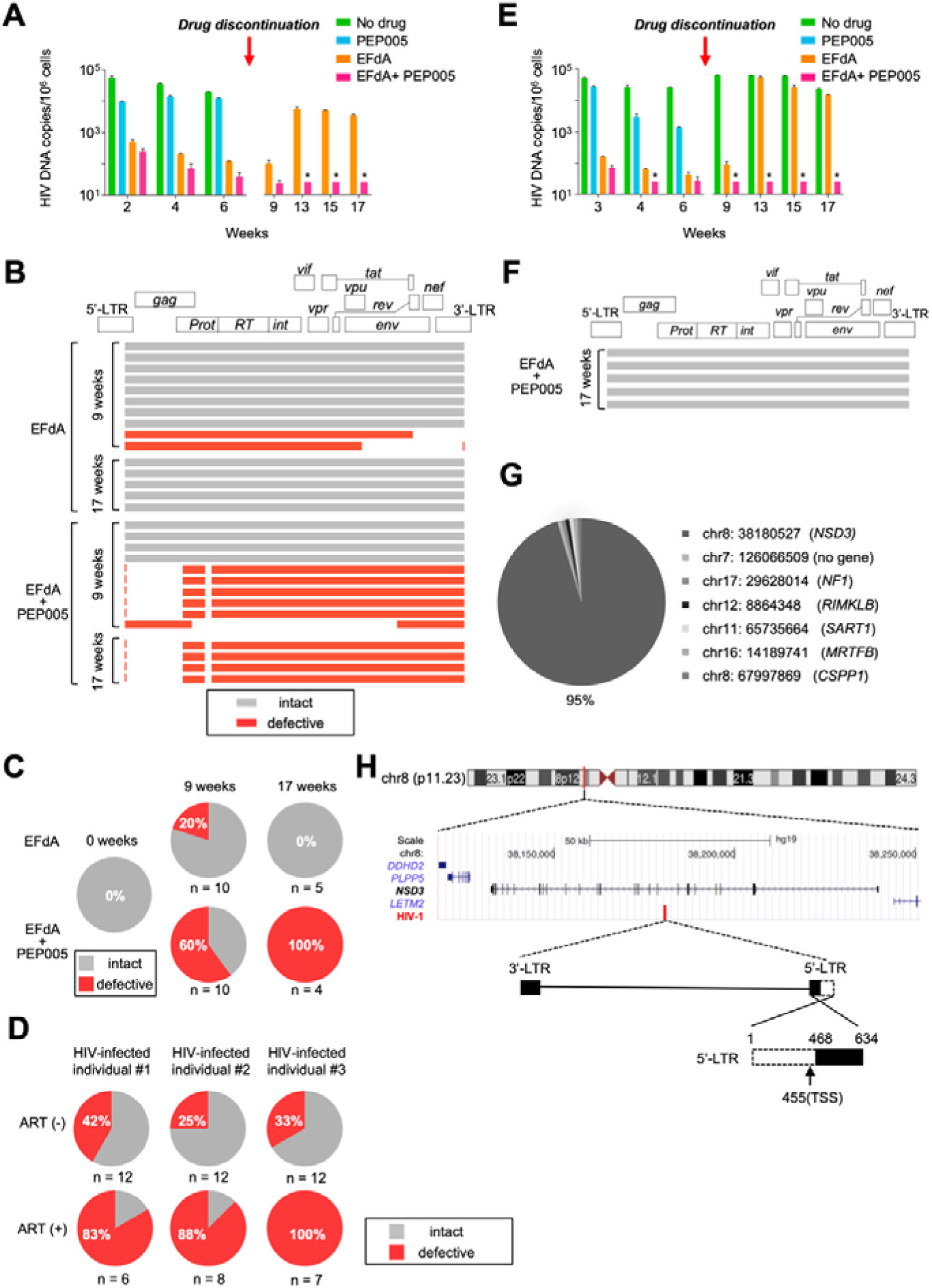
Mechanisms for the elimination of HIV-1 producible cells *in vitro*. **(A)** Quantification of intracellular copies of HIV-1 DNA at each time point in Exp. 6 (fig. S3A). **(B)** Schematic representation of the individual provirus structures from two different treatment groups and at two time points in Exp. 6. Each horizontal bar represents an individual HIV-1 genome, as determined by amplification of near full-length HIV-1 DNA from a single HIV-1 genome and DNA sequencing. The gray bars denote full-length types and the red bars indicate defective proviruses. **(C)** Pie charts reflecting the proportion of defective and intact proviruses in Exp. 6. **(D)** Pie charts reflecting the proportion of defective and intact proviruses in PBMCs from three HIV-infected individuals. (**E**) Quantification of intracellular copies of HIV-1 DNA at each time point in Exp. 1 (Fig. 2D). **(F)** Schematic representation of the individual provirus structures in Exp. 1 for the EFdA/PEP005 culture group 17 weeks after drug treatment initiation. **(G)** Pie chart showing the relative abundance of each HIV-1-infected clone. Chromosomal number and position of each clone is shown in the right panel. (**H)** Schematic figure of the provirus structure and IS in the expanded clone. A 467-bp deletion in the 5′-end of 5′LTR was observed. TSS, transcription start site. Asterisk (*) stands for below detection limit.

We also analyzed HIV-1 proviruses in Exp.1 (Fig. 2D) and similarly found that the HIV-1 DNA load was markedly decreased in samples treated with EFdA alone or EFdA together with PEP005 (Fig. 5E). Nearly full-length HIV-1 PCR and sequencing analysis (*34*) revealed that all remaining proviruses under combination treatment did not contain any critical mutations or deletions in the regions coding for viral proteins (Fig. 5F). However, these cells did not transcribe HIV-1 mRNA after TNF-α stimulation (fig. S3B), leading us to make the hypothesis that there might be some deletions or mutations in the provirus outside the primer-binding sites of nearly full-length HIV-1 PCR. To test the hypothesis, we analyzed the whole sequence of the 5’LTR. First, we determined HIV-1 ISs by LM-PCR (*26*) to design PCR primers to amplify the junction between the host and the viral genome. Since we can obtain the information of clonal abundance of each HIV-1-infected clone by LM-PCR (*10, 11, 26*), we incidentally found that one infected clone was remarkably expanded (Fig. 5G). A part of the 5′LTR, including the transcription start site, in the expanded clone was actually deleted (Fig. 5H and fig. S3C). This can explain the observed lack of virus rebound. Interestingly, *NSD3* gene were known as a cancer-related gene targeted by HIV-1 integration in an HIV-1-infected individual under cART (*10*).

In this study, we observed the following mechanisms underlying successful elimination of HIV-1 producible cells *in vitro*: (1) large deletion(s) in viral protein-coding regions; (2) critical mutation(s), such as nonsense mutations and frame-shift mutations (fig. S5), in the viral coding sequence; and (3) abnormalities in HIV-1 proviral transcription (a schematic diagram is shown in fig. S6) (*21, 34, 35*). A long-term antiretroviral treatment in HIV-1-infected individuals, reduces the numbers of replication-incompetent proviruses and increased the proportion of defective HIV-1 proviruses in a previous (*21*) and current study (Fig5. D and fig. S4 A and B). However, it is likely that a minor cell population with replication-competent HIV-1 proviruses persists during the long period of cART, maintaining the ability to reverse HIV-1 latency (*8, 14*). We demonstrated that the addition of an LRA to antiretroviral drug seemed to accelerate the elimination of cells infected with replication-competent HIV-1 proviruses that exist even as a minor population in Jurkat/NL cells.

## DISCUSSION

The usage of LRAs aims to reactivate latent proviruses to induce production of viral antigens and/or HIV-1 virions. A number of studies have demonstrated that recently developed small-molecule compounds have the ability to reverse latently HIV-1 infected cells (*6, 7*). However, Battivelli et al (*36*) reported that LRAs could reactivate only a part of infected cells with latent proviruses using their *in vitro* model, suggesting that there is a wide variation in drug susceptibility to LRAs among different HIV-1-infected clones. Genetic and epigenetic environment of integrated provirus plays a role in such different drug susceptibility. HIV-1 preferentially integrates into the genomic region with active transcription, resulting in a high proportion of HIV-1 integration within the host gene body (*27*) (Fig. 1F). Transcriptional interference between the host genes and integrated proviruses is another factor that affects proviral transcription (*37, 38*). In line with this notion, a recent study reported that there is a higher proportion of intact proviruses integrated in the opposite orientation relative to the host genes in CD4^+^ T cells of HIV-1-infected individuals under cART (*13*). In addition, epigenetic status of the integrated provirus is associated with accessibility of transcription factors that drive promoter activity of the 5_LTR. These findings indicate that susceptibility to LRAs among different HIV-1 clones is variable depending on the genetic and epigenetic environments of integrated proviruses. Thus, we need to develop drug combination that is effective to reactive a wide variety of HIV-1 provirus to eliminate cells with replication-competent provirus. However, currently available *in vitro* latent models to evaluate LRA activity carry only one or two integrated HIV-1 proviruses (Fig. 1D). Thus, a compound can potently reactivate a specific HIV-1 provirus in a latent cell line but may not do so for other clones. To our knowledge, this is the first report describing *in vitro* model available for evaluating the long-term effect of LRAs on a variety of HIV-1 clones. Here we show that WIPE assay enables us to analyze the effect of drug treatment on several thousand different clones with a similar distribution of HIV-1 proviruses as observed *in vivo* (Fig. 1E and F). The WIPE assay along with an in-depth characterization of the HIV-1 provirus would provide further insights on the underlying mechanism of HIV-1 latency, which cannot be obtained using conventional latent cell lines. Further studies using the WIPE assay may also provide mechanistic insights on molecular targeting, not only by LRAs but also by novel strategies; for example, the “block-and-lock” strategy was recently proposed, in which particular agents lock the HIV-1 promoter in a deep latency state to prevent viral reactivation (*39, 40*).

We utilized the WIPE assay to evaluate the reduction and eradication of replication competent HIV-1-DNA after a combination therapy of existing ART drug(s) with LRA(s). In the Jurkat/NL system, there is a continuous and dynamic viral infection, including *de novo* infection, cell apoptosis triggered by viral production, replication of uninfected cells, and generation of latently infected cells. ART drugs inhibit *de novo* infection from infected to uninfected cells, while LRAs activate latently infected cells and induce reactivation of viral antigen expression. As there are no antiviral CTLs and antibodies in the WIPE assay, elimination of reactivated cells is mostly due to viral cytopathicity or apoptosis of the reactivated cells. Notably, we recently reported that some LRAs, such as PEP005, strongly induce the upregulation of active caspase-3, resulting in enhanced apoptosis (*32*). That may explain why the addition of an LRA successfully accelerated the elimination of latently HIV-1-infected cells in the WIPE assay even in the absence of the host immune system. In fact, our quantitative analysis with the mathematical model revealed that PEP005-treatment reduced 52.2% of the time until reactivation in WIPE assay (Fig. 4B).

From virological point of view, we used the CXCR4-tropic (X4-tropic) HIV-1 variant NL4-3 to infect a T cell-derived Jurkat cell line. There is emerging evidence showing the importance of HIV-1-latency in monocytes or macrophages with the CCR5-tropic (R5-tropic) HIV-1 variant (*41*). Thus, we obtained PM1CCR5 cells infected with R5-tropic HIV-1_JRFL_ (PM1CCR5/JRFL) but failed to maintain long-term chronic infection to evaluate drug efficacy (data not shown). For future research, cell culture models with R5-tropic HIV-1 infected, monocyte-derived cells would be important for analyzing the differences in the latency mechanisms between the X4-tropic and R5-tropic HIV-1 variants.

We use a CD4^+^ T-cell line, Jurkat cells, as host cells in the WIPE assay, but there should be a wide variation of the host CD4^+^ T cell *in vivo* in infected individuals. That also can explain different susceptibility of HIV-1-infected CD4^+^ T cell against various LRAs (*36*). Therefore, we propose the use of the WIPE assay to evaluate candidate LRA drugs for initial evaluation. Then, compounds with potent activity in the WIPE assay can be further evaluated by long-term drug assays using primary CD4^+^ T cell-derived HIV-1 latent reservoir models (*42*). or animal models (i.e., HIV-1-infected humanized mice or SIV-infected macaques) (*22-24*). This strategy may facilitate an increase in the efficiency of drug development for LRAs and help to identify potent LRAs to reduce the reservoir size *in vivo*.

Taken together, our findings provide a proof-of-concept for the shock and kill strategy against HIV-1 infection using our newly established *in vitro* assay. A combination of the persistent and heterogeneous HIV-1 infection *in vitro* model and high-throughput characterization of HIV-1 proviruses will be useful in developing a new generation of LRAs specific for HIV-1 proviral latency and for optimizing drug combinations to reduce the HIV-1 latently infected cells.

## MATERIALS AND METHODS

### Drugs and reagents

The anti-HIV-1 reverse-transcriptase inhibitor EFdA/MK-8591/ISL and the protease inhibitor DRV were synthesized, as previously described (*29*). PEP005 (PKC activator) was purchased from Cayman Chemical (Ann Arbor, MI); SAHA (vorinostat; HDAC inhibitor) from Santa Cruz Biotechnology (Dallas, TX); JQ-1 (BRD4 inhibitor) from BioVision (Milpitas, CA); GSK525762A (BRD4 inhibitor) from ChemScene (Monmouth Junction, NJ); Ro5-3335 from Merck (Darmstadt, Germany); and Al-10-49 (CBFβ/RUNX inhibitor) from Selleck (Houston, TX). Prostratin and Bryostatin-1 (PKC activator) were purchased from Sigma-Aldrich (St. Louis, MO), while Panobinostat (HDAC inhibitor), GS-9620 (TLR-7 agonist), and Birinapant (IAP inhibitor) were purchased from MedChem Express (Monmouth Junction, NJ). Phorbole 12-myristate13-acetate (PMA) and TNF-α were purchased from Wako Pure Chemical (Osaka, Japan) and BioLegend (San Diego, CA), respectively.

### Establishment of the HIV-1 chronically infected cell culture model

Various T cell-derived cell lines [Jurkat, MT-4, Hut78, MOLT4 (ATCC), Jurkat-LTR-GFP (JLTRG), and PM1-CCR5 (NIH AIDS Reagent Program)] were used to obtain cell populations with chronic HIV-1 infection. Cells were infected with HIV-1_NL4-3_ or HIV-1_JRFL_ (PM1-CCR5) and cultured in RPMI 1640 medium (Sigma-Aldrich) supplemented with 10% fetal calf serum (Sigma-Aldrich), 50 U/mL penicillin, and 50 μg/mL kanamycin. Cells were passaged weekly to maintain cell numbers < 5 × 10^6^ cells/mL when confluent. p24 levels in the supernatant were monitored using Lumipulse G1200 (FUJIREBIO, Tokyo, Japan). The number of cells with intracellular p24 was also monitored on day 30 after infection by flow cytometry (as described below).

### HIV-1 reversal in latently infected cells and caspase-3 activation by LRAs

Chronically HIV-1_NL4-3_-infected Jurkat/NL cells were treated with 1 of the 11 drugs (1 μM) for 24 h, after which changes in supernatant p24 levels and induction of caspase-3 activation were determined by flow cytometry.

### Flow cytometry analysis

The ratios of intracellular HIV-1 p24^+^ cells, GFP^+^ cells, and the active form of caspase-3 expression were determined as previously described (*8, 32*). Briefly, Jurkat/NL or JLTRG/NL cells were washed twice with phosphate buffered salts (PBS) and stained with Ghost Dye Red 780 (TONBO Biosciences, San Diego, CA) for 30 min at 4°C. The cells were then fixed with 1% paraformaldehyde/PBS for 20 min, and permeabilized in a flow cytometry perm buffer (TONBO Biosciences). After 5-min incubation at room temperature (25–30°C), cells were stained with anti-HIV-1 p24 (24-4)-fluorescein isothiocyanate (FITC) monoclonal antibody (mAb; Santa Cruz Biotechnology) and/or Alexafluor 647-conjugated anti-active caspase-3 mAb (C92-605; BD Pharmingen, San Diego, CA) for 30 min on ice. For propidium iodide (PI)/annexin V staining, cells were washed twice with PBS and resuspended in annexin V binding buffer (BioLegend) at a concentration of 1 × 10^7^ cells/mL. The cells were then stained with FITC annexin V (BioLegend) and PI solution (BioLegend) for 15 min at room temperature. Cells were analyzed using a BD FACSVerse flow cytometer (BD Biosciences, Franklin Lakes, NJ). Data collected were analyzed using FlowJo software (Tree Star, Inc., Ashland, OR).

### Sorting of GFP^+^ or GFP^-^ cells from HIV-1_NL4-3_-infected JLTRG cells

HIV-infected Jurkat-LTR-GFP cells (6□×□10^6^) were resuspended in FACS buffer (PBS with 1% fetal calf serum), after which GFP^+^ or GFP^−^ cells were sorted using BD FACS Aria I (BD Biosciences). The sorted GFP^−^ cells were stimulated with 10 ng/mL TNF-α for 6 h and then GFP expression levels were analyzed using BD FACSVerse (BD Biosciences). The level of *gag* expression after 18-h stimulation with 10 ng/mL TNF-α was analyzed by reverse-transcription (RT)-digital droplet PCR (ddPCR). ddPCR droplets were generated using the QX200 droplet generator (Bio-Rad Laboratories, Hercules, CA). RT-PCR was performed using a C1000 Touch thermal cycler (Bio-Rad Laboratories) with the primers listed in table S2. The *gag*-positive and negative droplets were quantified based on fluorescence using the QX200 droplet reader (Bio-Rad Laboratories).

### Determination of antiviral activity of LRAs and conventional anti-HIV-1 drugs in Jurkat/NL cells (WIPE assay)

Jurkat/NL cells (5.0 × 10^4^ cells/ml) were treated with a drug (e.g., EFdA, DRV, or PEP005) or a combination of drugs in a 12-well plate. Culture medium was exchanged and the drug was added. Drug treatment was stopped approximately on week 9 and the culture was maintained for an additional 8 weeks without drug supplementation. Supernatant p24 levels and intracellular HIV-1-DNA levels were monitored weekly during cell culture. At the end of each experiment, drug-treated cells with low/undetectable supernatant p24 levels were stimulated with 10 ng/mL TNF-α to confirm viral recurrence.

### Isolation of PBMCs from HIV-1 infected individuals

The study was performed in accordance with the guidelines of the Declaration of Helsinki. Analysis of clinical samples shown in Fig. 1E was conducted based on a protocol reviewed and approved by the Kumamoto University (Kumamoto, Japan) Institutional Review Board (approval number Genome 258).

Peripheral blood samples analyzed as shown in fig. S4 were collected from individuals infected with HIV-1 before or after receiving cART for at least 7 years. The Ethics Committee of the National Center for Global Health and Medicine (Tokyo, Japan) approved this study (NCGM-G-002259-00). Informed written consent was obtained from all participants (table S1) prior to the study. All subjects maintained low viral loads (< 20 copies/mL; except for occasional blips) during therapy. CD4^+^ T-cell counts in peripheral blood samples ranged from 447 to 632 cells/mm^3^ (average 529 cells/mm^3^). The plasma viral loads were < 20 copies/mL, as determined by quantitative PCR (Roche COBAS AmpliPrep/COBAS TaqMan HIV-1 Test version 2.0) at the time of enrollment in the study. PBMCs were isolated from whole blood by density-gradient centrifugation using Ficoll-Paque™ (GE Healthcare, Chicago, IL). Total cellular DNA was extracted and used in subsequent PCR experiments.

### Quantification of intracellular HIV-1 DNA levels

Total cellular DNA was extracted from cells (cell lines or PBMCs) using a QIAmp DNA Blood mini kit (Qiagen, Hilden, Germany) according to manufacturer’s instructions. Quantitative PCR (qPCR) analysis of intracellular HIV-1 DNA levels was conducted using Premix Ex Taq (Probe qPCR) Rox plus (Takara Bio, Kusatsu, Japan). The oligonucleotides HIV-1 LTR and β2-microglobulin were used for HIV-1 DNA quantification and cell number determination, respectively (primer sequences are provided in table S2 (*26, 43, 44*). HIV-1 proviral DNA copy and cell numbers were calculated based on a standard curve generated using a serially diluted pNL4-3 plasmid and DNA extracted from Jurkat cells, respectively.

### RT-qPCR for HIV-1 mRNA quantification

Total cellular RNA was extracted from Jurkat cells infected with HIV-1 using the RNeasy Mini Kit (Qiagen) according to manufacturer’s instructions. cDNA was synthesized using the PrimeScript RT Master Mix (Takara Bio). RT-qPCR analysis of intracellular HIV-1 RNA was performed using PowerUp SYBR Green Master Mix (Applied Biosystems, Foster City, CA). Primer sequences used for the detection of HIV-1-RNA and β-actin gene are listed in table S2. To determine the reactivation of HIV-1 in Jurkat/NL cells, relative HIV-1-RNA expression levels were normalized to that of β-actin gene.

### Amplification of near-full-length single HIV-1 genome and sequencing

Nearly full-length single HIV-1 genome PCR was performed as described previously (*34*) with minor modifications. Briefly, genomic DNA was diluted to the single-genome level based on ddPCR and Poisson distribution statistics. The resulting single genome was amplified using Takara Ex Taq hot start version (first-round amplification). PCR conditions for first-round amplification consisted of 95°C for 2 min; followed by 5 cycles of 95°C for 10 s, 66°C for 10 s, and 68°C for 7 min; 5 cycles of 95°C for 10s, 63°C for 10 s, and 68°C for 7 min; 5 cycles of 95°C for 10 s, 61°C for 10 s, and 68°C for 7 min; 15 cycles of 95°C for 10 s, 58°C for 10 s, and 68°C for 7 min; and finally, 68°C for 5 min. First-round PCR products were diluted 1:50 in PCR-grade water and 5 μL of the diluted mixture was subjected to second-round amplification. PCR conditions for the second-round amplification were as follows, 95°C for 2 min; followed by 8 cycles of 95°C for 10 s, 68°C for 10 s, and 68°C for 7 min; 12 cycles of 95°C for 10 s, 65°C for 10 s, and 68°C for 7 min; and finally, 68°C for 5 min. Primer information is provided in table S2. PCR products were then visualized by electrophoresis on a 1% agarose gel. Based on Poisson distribution, samples with ≤ 30% positive reactions were considered to contain a single HIV-1 genome and were selected for sequencing. Amplified PCR products of the selected samples were purified using a QIAquick PCR Purification Kit (Qiagen) according to manufacturer’s instructions. Purified PCR products were sheared by sonication using a Picoruptor device (Diagenode, Liege, Belgium) to obtain fragments with an average size of 300–400 bp. Libraries for next-generation sequencing (NGS) were prepared using the NEBNext Ultra II DNA Library Prep Kit for Illumina (New England Biolabs, Ipwich, MA) according to manufacturer’s instructions. Concentration of library DNA was determined using the Qubit dsDNA High Sensitivity Assay kit (Invitrogen, Carlsbad, CA). The libraries were subsequently pooled together, followed by quantification using the Agilent 2200 TapeStation and quantitative PCR (GenNext NGS Library Quantification kit; Toyobo, Osaka, Japan), and sequenced using the Illumina MiSeq platform (Illumina, San Diego, CA). The resulting short reads were cleaned using an in-house Perl script (kindly provided by Dr. Michi Miura, Imperial College London, UK), which extracts reads with a high index-read sequencing quality (Phred score > 20) in each position of an 8-bp index read. Next, adapter sequences from Read1 and Read2 were removed, followed by a cleaning step to remove reads that were too short or had a very low Phred score, as previously described (*26*). The clean sequencing reads were aligned with the NL4-3 reference genome (GenBank-M19921) using the BWA-MEM algorithm (*45*). Further data processing and cleanup, including the removal of reads with multiple alignments and duplicated reads, were performed using Samtools (*45*) and Picard (http://broadinstitute.github.io/picard/). The aligned reads were visualized using Integrative Genomics Viewer (*46*), and consensus sequences were copied and aligned using MUSCLE (*47*). NGS analyses of nearly full-length HIV-1 PCR products from PBMCs of HIV-1-infected individuals were conducted using MinION platform with Flow Cell R9.4.1 and Rapid Barcoding kit (Oxford Nanopore Technologies, Oxford, UK), according to manufacturer’s instructions. Sequencing reads cleaned using EPI2ME software (Oxford Nanopore Technologies) were aligned and analyzed as described above.

### Ligation-mediated PCR (LM-PCR)

Detection of HIV-1 ISs was performed using ligation-mediated PCR and high-throughput sequencing, as previously described (*26*) but with minor modifications. Briefly, cellular genomic DNA was sheared by sonication using the Picoruptor device to obtain fragments with an average size of 300–400 bp. DNA ends were repaired using the NEBNext Ultra II End Repair Kit (New England Biolabs) and a DNA linker (*26*) was added. The junction between the 3′LTR of HIV-1 and host genomic DNA was amplified using a primer targeting the 3′LTR and a primer targeting the linker (*26*). PCR amplicons were purified using the QIAquick PCR Purification Kit (Qiagen) according to manufacturer’s instructions. This was followed by Ampure XP bead purification(Beckman Coulter). Purified PCR amplicons were quantified using Agilent 2200 TapeStation and quantitative PCR (GenNext NGS library quantification kit; Toyobo). LM-PCR libraries were sequenced using the Illumina MiSeq as paired-end reads, and the resulting FASTQ files were analyzed as previously described (*26*). A circos plot showing virus ISs in the Jurkat/NL4-3 model and different cell lines was constructed using the OmicCircos tool available as a package in R software (*48*).

### Bioinformatic analysis

Bed files containing the IS information were generated from the analyzed exported files. Data on RefSeq genes were obtained using UCSC Genome Browser (https://genome.ucsc.edu/) and the positions of RefSeq genes were compared with those of IS using the R package hiAnnotator (http://github.com/malnirav/hiAnnotator).

### Statistical analysis

Differences between groups were analyzed for statistical significance using the Mann-Whitney U test and log-rank test. Data were analyzed using a chi-squared test with Prism 7 software (GraphPad Software, Inc., La Jolla, CA), unless otherwise stated. Statistical significance was defined as *P* < 0.05.

## SUPPLEMENTARY MATERIALS

Figs. S1 to S8

Tables S1 to S5

Supplementary Text for mathematical model and statistical analysis

Supplementary references

## Acknowledgements

We thank Jumpei Ito for providing access to the Python program, allowing us to perform association analyses between the epigenetic environment and viral integration sites. We also thank Shinichi Oka for his advice and suggestions for the manuscript.

## Author contributions

K.Y., Y.S., and Ke.M. designed the study. Ko.M., Sa.I., K.T., S.H., H.K., M.M., and N.S.D. performed the experiments. S.M. provided clinical sample DNA. Sa.I., B.J.Y.T., and Y.S. performed bioinformatic analysis. T.T., K.K., K.S.K., and Sh.I. performed mathematical analysis. H.G., S.O., S.M., H.M., Y.S., and Ke.M. supervised the work. Y.S. and Ke.M. wrote the manuscript with input from all authors.

