## Supplementary Materials for "A widely-distributed HIV-1 provirus elimination assay to evaluate latency-reversing agents *in vitro*"

**This PDF file includes:**

- Figs. S1 to S8
- Tables S1 to S5
- Supplementary Text for mathematical model and statistical analysis
- Supplementary references

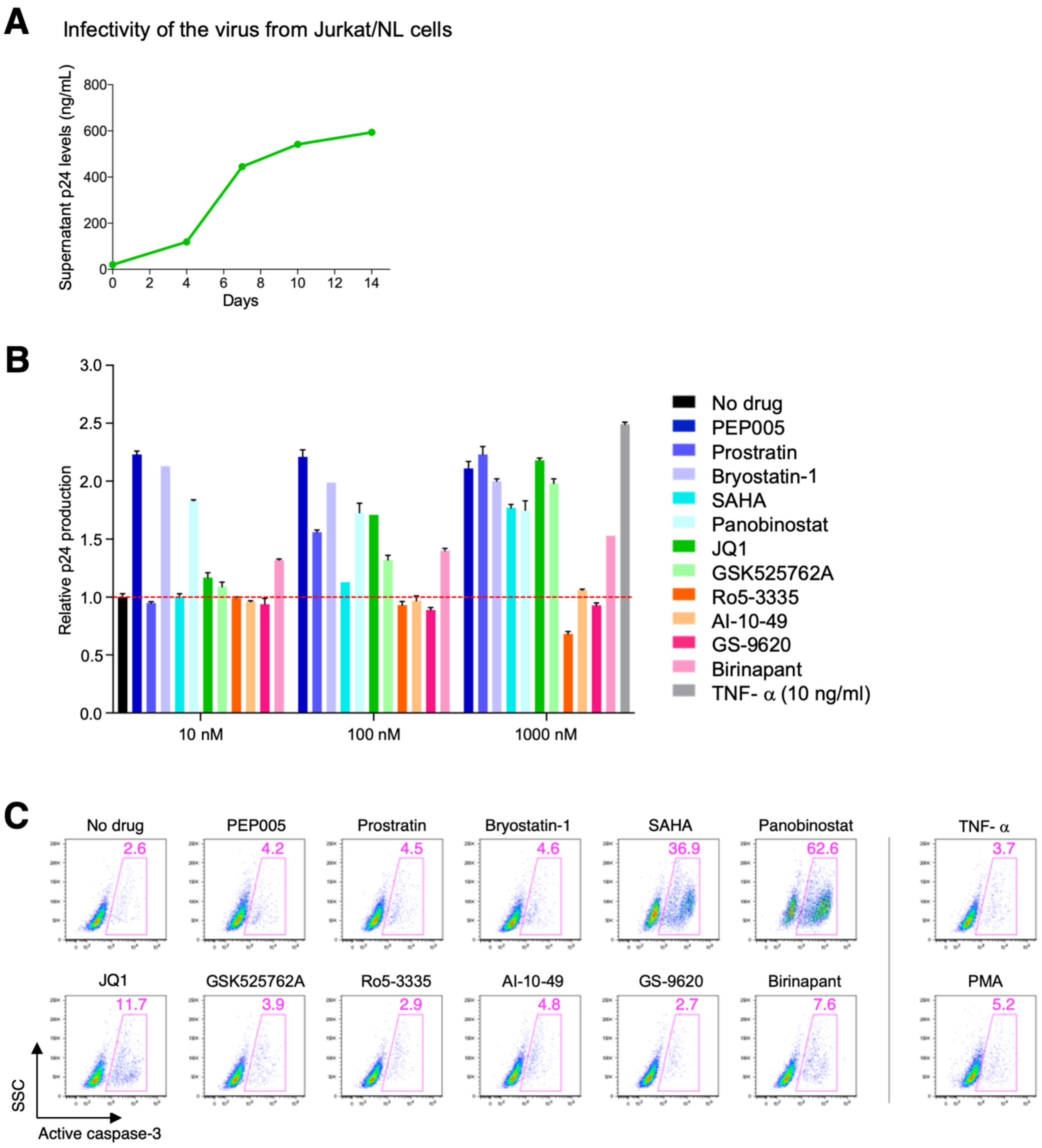


**fig. S1. Viral infectivity in Jurkat/NL cells and effects of LRAs on HIV-1 production and cell apoptosis.** **(A)** Infectivity of HIV-1$_{NL4-3}$ produced from Jurkat/NL cells. MT4 cells were infected with the virus, cultured, and then supernatant p24 levels were measured. (**B**–**C)** Efficacy of LRAs in inducing HIV-1 production or caspase-3 activation in Jurkat/NL cells. Cells were treated with a drug (1 μM) for 24 h and the changes in supernatant p24 values (**B**) or percentage of active forms of caspase-3 expression (**C**) were examined.

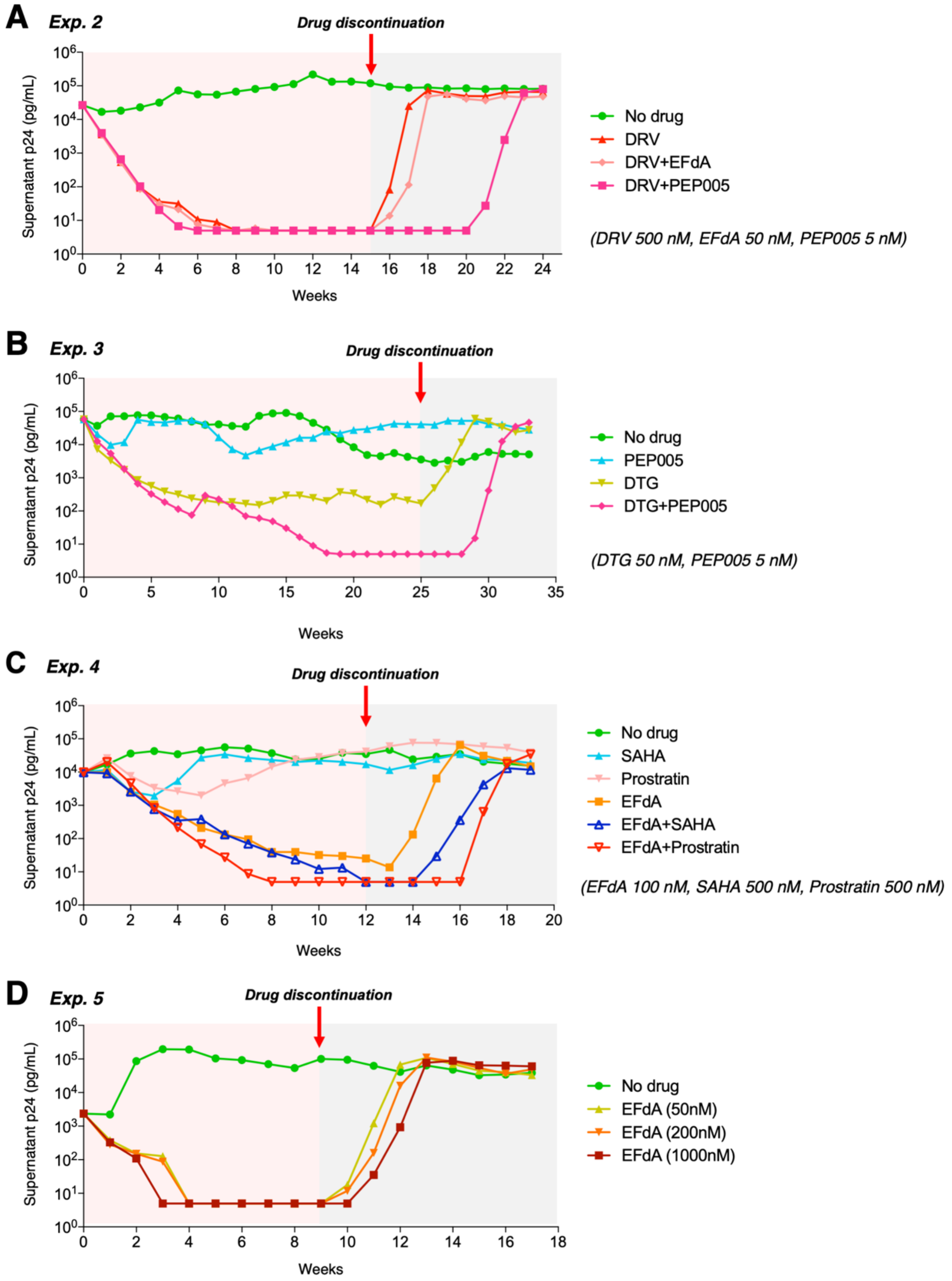


**fig. S2. Effect of various combinations of antiretroviral drugs and LRAs on viral persistence.** Changes in HIV-1 production under treatment with 5 nM PEP005, 50 nM EFdA, and/or 500 nM Darunavir (DRV, protease inhibitor) (**A**), 5 nM PEP005, and/or 50 nM Dolutegravir (DTG, integrase inhibitor) (**B**), and 100 nM EFdA, 500 nM SAHA (HDAC inhibitor), and/or 500 nM prostratin (PKC activator) (**C**). **(D)** EFdA at different concentrations (50 nM, 200 nM, and 1 µM) was examined. A higher concentration of EFdA (200 nM and 1 µM) slightly delayed the recurrence of supernatant viruses after treatment interruption.

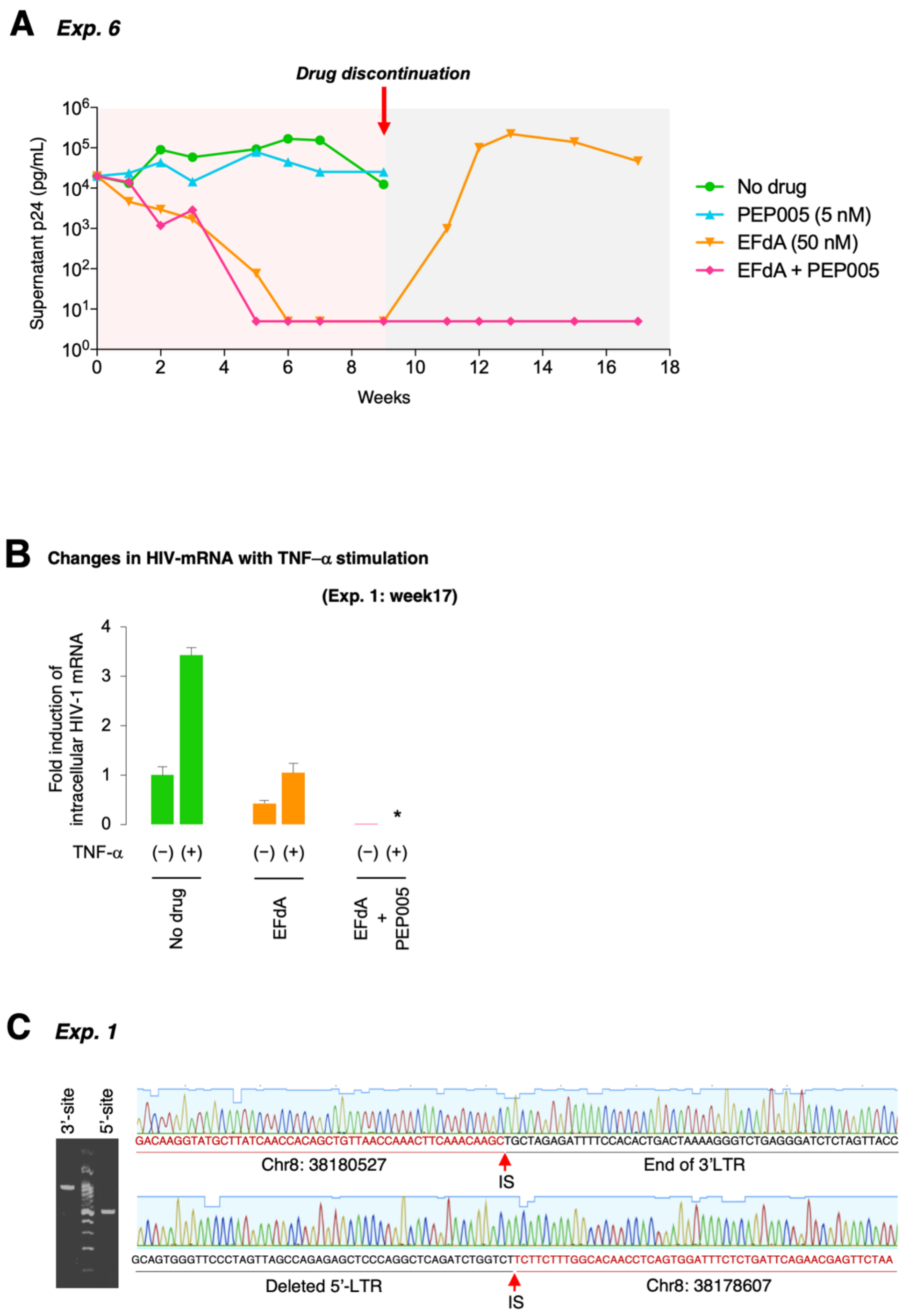


**fig. S3. Underlying mechanisms of the experimental cure (Exp. 1 and 6). (A)** Result of WIPE assay with EFdA and PEP005 in Exp. 6 (an experiment shown in Fig. 2B). **(B)** Analysis of HIV-1 mRNA transcripts in cells of Exp. 1 (Fig. 2D) on week 17 with TNF-α stimulation. Cells were treated with 10 ng/mL TNF-α for 24 h, and the change in intracellular HIV-1-mRNA transcripts was analyzed. **(C)** Result of IS-specific PCR of the expanded clone in Exp. 1 (Fig. 3F). PCR bands amplified from either 5′LTR- or 3′LTR-host junctions (502 bp and 773 bp, respectively) are shown on the left. DNA sequencing results of the host-virus junctions are shown on the right.

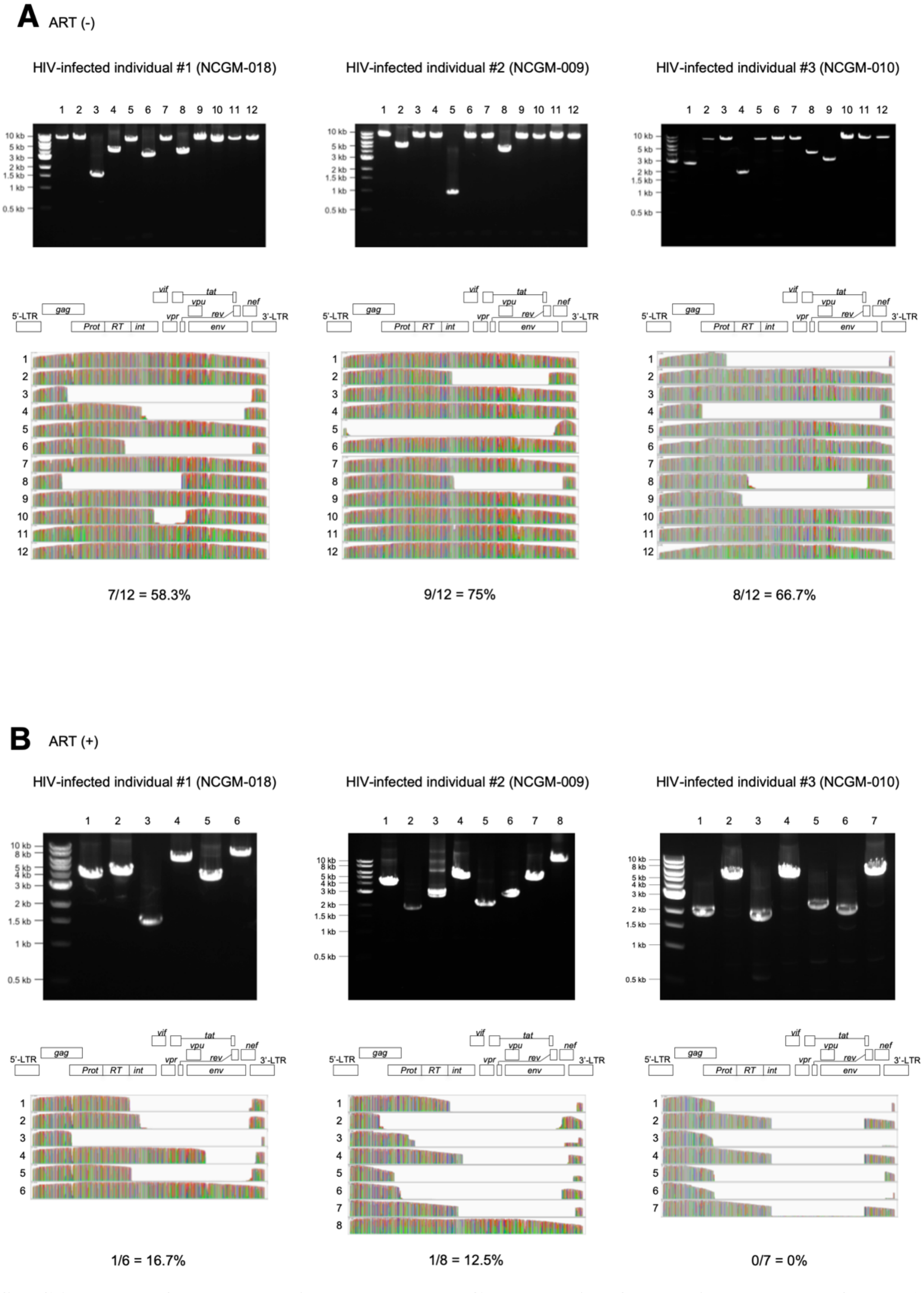


**fig. S4. Nearly full-length, single-genome PCR analysis of the primary cells of HIV-1 infected patients.** **(A)** PCR products of cells from three HIV-1 patients (table S2) before initiation of effective cART treatment. **(B)** PCR products of cells from the same patients after cART treatment (duration, 84–264 months).

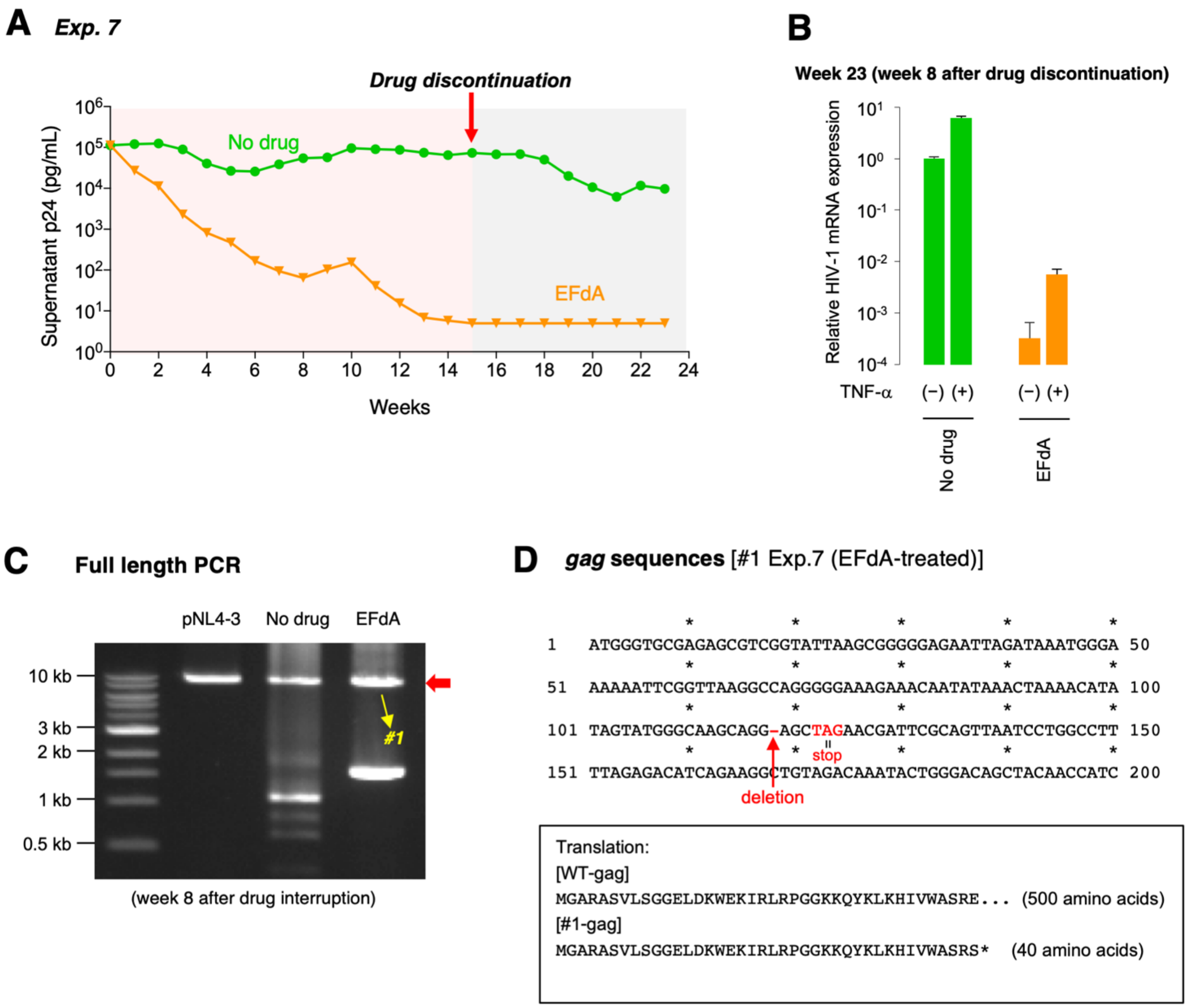


**fig. S5. An underlying mechanism of an experimental cure (Exp. 7). (A)** Treatment of Jurkat/NL cells with EFdA (Exp. 7). In this experiment, HIV-1 rebound was not observed in cells treated with EFdA until week 23; however, EFdA-treated cells showed an increase in HIV-1 mRNA expression with TNF-α stimulation (**B**), suggesting that the cells containing replication-competent proviruses are a minor population. **(C)** PCR products (9031 bp) of cell samples from Exp. 7 on week 23. **(D)** Sequencing analysis of PCR products (#1) in (**C**) with NGS demonstrated that there is a 1-bp deletion with a premature stop codon in HIV-1 *gag*.

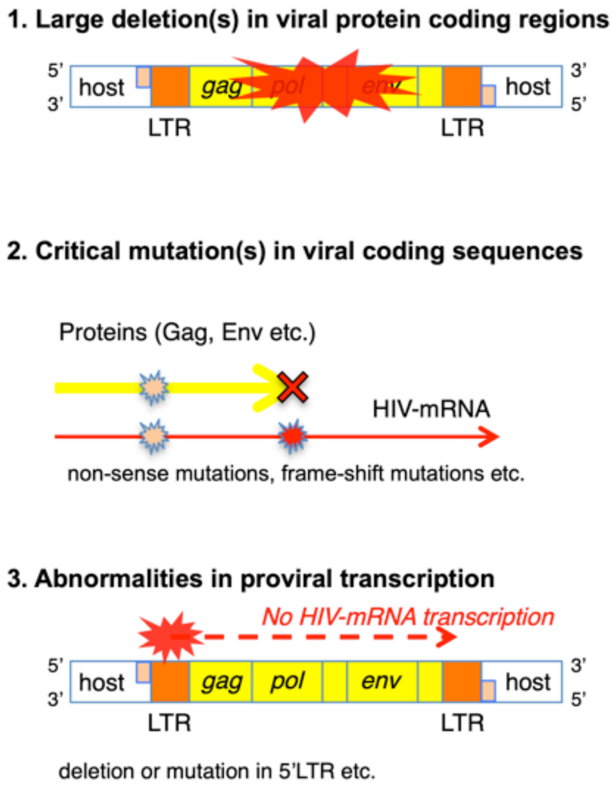


**fig. S6. Mechanism of HIV-1 provirus replication incompetency observed after drug treatment.**

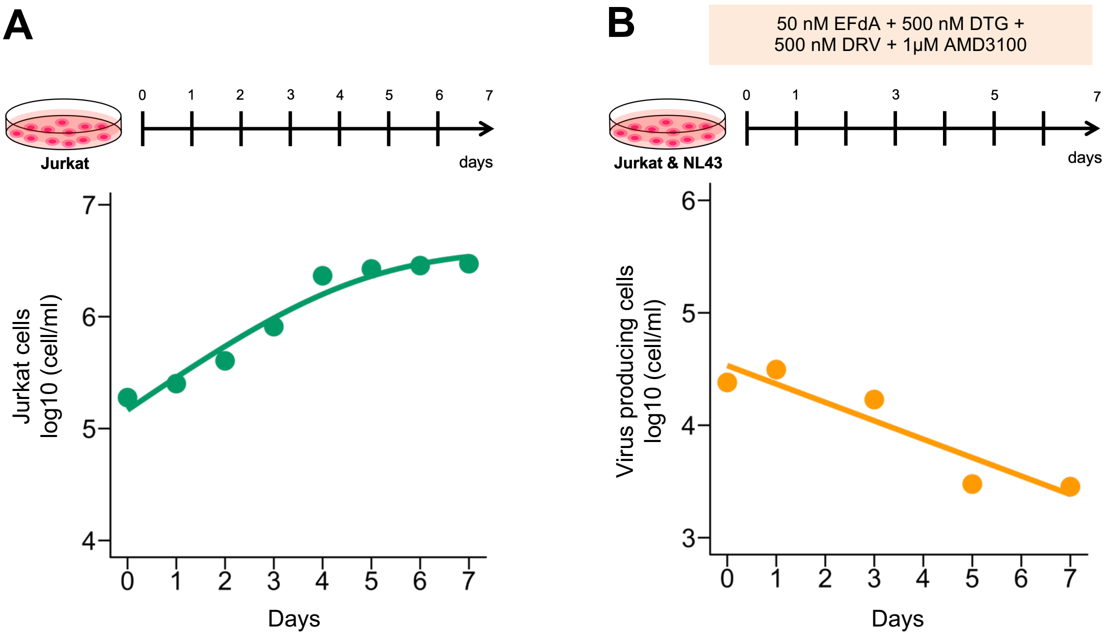


**fig. S7. Dynamics of jurkat cell growth and virus-producing cell death. (A)** By counting the cells for 7 days, the growth kinetics of jurkat cells was estimated as described in Supplementary text. **(B)** By counting the cells for 7 days in the presence of antiviral drugs, the death kinetics of virus-producing (p24-positive) infected cells was estimated as in Supplementary text.

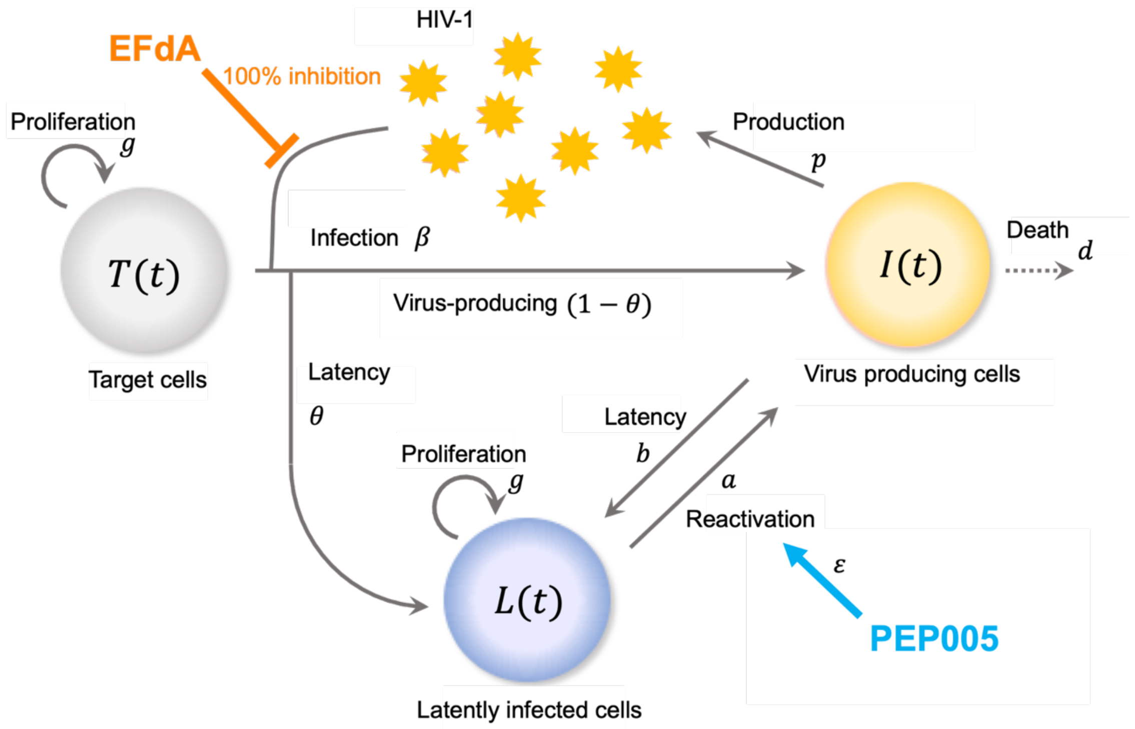


**fig. S8. Schematic representation of HIV-1 infection dynamics with antiviral drugs in WIPE assay.** The uninfected target cells, $T(t)$, are assumed to grow in a logistic manner as described in Supplementary text, and to be infected by viruses, $V(t)$, at rate $\beta$. It is assumed that a fraction $\theta$ becomes latently infected cells, $L(t)$, with the logistic growth, and $1-\theta$ is virus-producing infected cells, $I(t)$, having death rate $d$. The latently infected cells are reactivated at rate $a$, then (re-)start virus production at rate $p$, while the virus-producing cells become latently infected cell at rate $b$. We also assume EFdA completely inhibits *de novo* infection (100% inhibition rate, $\beta=0$) and PEP005 enhances the reactivation rate to $\varepsilon a$ where $\varepsilon>1$.

**table S1.** Characteristics of HIV-1-infected patients

| Patient ID | M/F | Age | VL[a] (copies/mL) | CD4 count[a] (cells/mm$^3$) | cART | Therapy (years) | Plasma HIV RNA < 20 copies/mL for (years) |
|---|---|---|---|---|---|---|---|
| NCGM-018 | F | 46 | <20 | 447 | FTC/TAF/EFV | 19 | 7 |
| NCGM-009 | M | 56 | <20 | 632 | FTC/TAF/RPV | 22 | 7 |
| NCGM-010 | M | 49 | <20 | 509 | FTC/TAF/COBI/EVG | 7 | 6 |

[a]VL and CD4 count were measured at the time of the study.

COBI, cobicistat; EFV, efavirenz; EVG, elvitegravir; FTC, emtricitabine; RPV, rilpivirine; TAF, tenofovir alafenamide fumarate; VL, viral load.

**table S2.** PCR primers used in the present study

[Quantitative PCR]

| Target | Primer name | Sequence (5′-3′) | Reference |
|---|---|---|---|
| HIV-1 LTR | MH531 (forward) | TGTGTGCCCGTCTGTTGTGT | (*1*) |
| | MH532 (reverse) | GAGTCCTGCGTCGAGAGAGC | (*1*) |
| | LRTp (probe) | FAM-CAGTGGCGCCCGAACAGGGA-BHQ1 | (*1*) |
| β2-microglobulin | β2m_S (forward) | GGAATTGATTTGGGAGAGCATC | (*2*) |
| | β2m_AS (reverse) | CAGGTCCTGGCTCTACAATTTACTAA | (*2*) |
| | β2m_P (probe) | FAM-AGTGTGACTGGGCAGATCATCCACCTTC-BHQ1 | (*2*) |
| HIV-1 gag | gag_S (forward) | GGTGCGAGAGCGTCGGTATTAAG | (*3*) |
| | gag _AS (reverse) | AGCTCCCTGCTTGCCCATA | (*3*) |
| β-actin | β-actin_S (forward) | GCGAGAAGATGACCCAGATC | (*4*) |
| | β-actin_AS (reverse) | CCAGTGGTACGGCCAGAGG | (*4*) |

[Near full-length single HIV-1 genome PCR]

| Primer set | Name | HXB2 position | Length (bp) | Sequence (5′-3′) | Reference |
|---|---|---|---|---|---|
| First round | DNA F1 | 623–649 | 9064 | AAATCTCTAGCAGTGGCGCCCGAACAG- | (*5*) |
| | DNA R1 | 9652–9676 | | TGAGGGATCTCTAGTTACCAGAGTC | (*5*) |
| Second round | Nested F | 638–666 | 8985 | GCGCCCGAACAGGGACYTGAAARCGAAAG | (*5*) |
| | DNA R2 | 9603–9632 | | GCACTCAAGGCAAGCTTTATTGAGGCTTA | (*5*) |
| Second (clinical) | DNA F2 | 682–705 | 8951 | TCTCTCGACGCAGGACTCGGCTTG | (*5*) |
| | DNA R2 | 9603–9632 | | GCACTCAAGGCAAGCTTTATTGAGGCTTA | (*5*) |

[Linker-mediated PCR primers]

| Target | Name | Sequence (5′-3′) | Reference |
|---|---|---|---|
| | Long linker | TCATATAATGGGACGATCACAAGCAGAAGACGGCATACGAGA TNNNNNNNN CGGTCTCGGCATTC CTGCTGAACCGCTCTTCCGATCT | (*6*) |
| | Short linker | p-GA TCGGAAGAGCGAAAAAAAAAAAAA | (*6*) |
| 1st PCR | B3 | GCTTGCCTTGAGTGCTTCAAGTAGTGTG | (*6*) |
| | B4 | TCATGATCAATGGGACGATCA | (*6*) |
| 2nd PCR | P5B5 | AATGATACGGCGACCACCGAGATCTACACGTGCCCGTCTGTTG TGTGACTCTGG | (*6*) |
| | P7 | CAAGCAGAAGACGGCATACGAGAT | (*6*) |

[High- throughput sequencing primers]

| Target | Name | Sequence (5′-3′) | Reference |
|---|---|---|---|
| HIV-1 | Read1 | ATCCCTCAGACCCTTTTAGTCAGTGTGGAAAATCTC | (*6*) |
| Human genome | Read2 | CGGTCTCGGCATTCCTGCTGAACCGCTCTTCCGATCT | (*6*) |
| Adaptor barcode | Index1 | GATCGGAAGAGCGGTTCAGCAGGAATGCCGAGACCG | (*6*) |

**table S3.** Estimated parameter values of cell growth dynamics

| Parameter name | Symbol | Unit | Value |
|---|---|---|---|
| Proliferation rate of Jurkat cells | $g$ | $day^{-1}$ | 0.708 |
| Carrying capacity of Jurkat cells | $K$ | cell/ml | $4.10\times10^{6}$ |
| Death rate of virus producing cells | $d$ | $day^{-1}$ | 0.377 |

**table S4.** Estimated parameter values of HIV-1 infection dynamics in WIPE assay

| Parameter name | Symbol | Unit | Value | 95% CI |
|---|---|---|---|---|
| Infection rate | $\beta$ | (p24/ml day)$^{-1}$ | $7.18\times10^{-6}$ | $(0.21 - 1.99)\times10^{-5}$ |
| Latency rate in new infection | $\theta$ | - | $2.51\times10^{-5}$ | $1.60\times10^{-6} – 1.23\times10^{-4}$ |
| Production rate of total viral protein | $p$ | p24/cell day$^{-1}$ | $7.25\times10^{-2}$ | $(0.29 – 1.51)\times10^{-1}$ |
| Reactivation rate of latently infected cells | $a$ | day$^{-1}$ | $3.76\times10^{-2}$ | $(1.92 – 6.30)\times10^{-2}$ |
| Latency rate of virus producing cells | $b$ | day$^{-1}$ | $4.05\times10^{-6}$ | $6.82\times10^{-7} – 1.17\times10^{-5}$ |
| Promoting effect of PEP005 on reactivation | $\varepsilon$ | - | 2.31 | 1.05 – 5.34 |

**table S5.** Estimated initial values

| Variable | Symbol | Unit | Value | 95% CI |
|---|---|---|---|---|
| Initial number of target cells | $T(0)$ | cell/ml | $1.11\times10^{6}$ | $(0.36\text{-} 2.60)\times10^{6}$ |
| Initial number of virus producing cells | $I(0)$ | cell/ml | $2.37\times10^{4}$ | $1.84\times10^{3}$ - $1.03\times10^{5}$ |
| Initial number of latently infected cells | $L(0)$ | cell/ml | $4.39\times10^{4}$ | $8.74\times10^{3}$ – $1.36\times10^{5}$ |
| Initial amount of HIV-1 | $V(0)$ | p24/ml | $2.03\times10^{4}$ | $(0.21$ - $7.62)\times10^{4}$ |

### Supplementary Text

<u>Quantification of cell growth and death</u>

We estimated the growth kinetics of jurkat cells using the following mathematical model:

$$\frac{dT(t)}{dt} = gT(t)\left(1 - \frac{T(t)}{K}\right)$$

where the variables $T(t)$ represents the numbers of uninfected target cells at time $t$, and the parameters $g$ and $K$ represent the growth rate and the carrying capacity of the cell culture well, respectively. Note that latently infected cells have same growth kinetics because of no intracellular viral replications due to inactivation. Nonlinear least-squares regression was performed to the time-course numbers of jurkat cells. The fitted parameter values are listed in table S3 and the model behavior using these best-fit parameter estimates is presented together with the data in fig. S7A.

In addition, to estimate the death rate of virus-producing infected cells, treatments with a combination of EFdA (50 nM), DTG (500 nM), DRV (500 nM) and AMD3100 (1 μM) were started after virus infection reached at steady state, and then measured the time-course numbers of those cells for 7 days. Since the drug combination perfectly inhibits *de novo* infection, the decay kinetics of the virus-producing cells is described by the following model:

$$\frac{dI(t)}{dt} = -dI(t)$$

where the $I(t)$ represents the numbers of virus-producing cells at time $t$, and the parameter $d$ represents the death rate. The fitted parameter values are listed in table S3 and the model behavior using these best-fit parameter estimates is presented together with the data in fig. S7B.

<u>Mathematical model describing HIV-1 infection dynamics in WIPE assay</u>

We here introduce an extended mathematical model considering reactivation of latent HIV-1 reservoirs for analyzing experimental data in WIPE assay (*7-9*);

$$\frac{dT(t)}{dt} = gT(t)\left(1 - \frac{N(t)}{K}\right) - \beta T(t)V(t) - d_p T(t),$$

$$\frac{dI(t)}{dt} = (1-\theta)\beta T(t)V(t) + \varepsilon a L(t) - \left(b + d + d_p\right)I(t),$$

$$\frac{dL(t)}{dt} = \theta\beta T(t)V(t) + gL(t)\left(1 - \frac{N(t)}{K}\right) + bI(t) - \left(\varepsilon a + d_p\right)L(t),$$

$$\frac{dV(t)}{dt} = pI(t) - d_p V,$$

where $T(t)$ is the numbers of uninfected target cells, $I(t)$ and $L(t)$ are the numbers of virus-producing and latently infected cells per ml of a culture (i.e., the total cells are $N(t) = T(t) + I(t) + L(t)$), respectively, and $V(t)$ is the viral load measured by the amount of HIV-1 p24 per ml of culture supernatant. The uninfected and latently infected cells grow at a rate $g$ with the carrying capacity of $K$ (the maximum number of cells in the cell culture flask). It is assumed that a fraction $\theta$ becomes latently infected cells, and $1-\theta$ is virus-producing infected cells having death rate $d$, when HIV-1 infect the target cells at rate $\beta$. The latently infected cells are reactivated at rate $a$, then (re-)start virus production at rate $p$ (implying only intact proviral DNA is assumed), while the virus-producing cells become latently infected cell at rate $b$. We also assume EFdA completely inhibits *de novo* infection (100% inhibition rate, $\beta = 0$) and PEP005 enhances the reactivation rate to $\varepsilon a$ where $\varepsilon > 1$. Note that $d_p$ represent the removal of virus and of the cells due to the experimental passage, and we fixed $d_p = 0.428$. In our earlier works (*10, 11*), we have shown that the approximating punctual removal as a continuous exponential decay has minimal impact on the model parameters and provides an appropriate fit to the experimental data.

Data fitting and parameter estimation

The parameters $g$, $K$ and $d$ were separately estimated as we discussed above, and fixed at $0.708$ per day, $4.10 \times 10^6$ cells and $0.377$ per day in table S3, respectively. A statistical model adopted from Bayesian inference assumed that measurement error followed a normal distribution with mean zero and constant variance (error variance). The non-informative prior distributions with upper and lower limits were used as the prior distributions of parameters. The posterior predictive parameter distribution as an output of MCMC computation represented parameter variability. To analyze our cell culture experimental datasets, we simultaneously fitted $T(t) + L(t)$, $I(t)$, $V(t)$ and $\left(I(t) + L(t)\right)/\{I(2\text{ week}) + L(2\text{ week})\}$ to the time-course of numbers of uninfected and latently infected (i.e., p24-negative) cells (cells/ml), virus producing (i.e., p24-positive) cells (cells/ml), supernatant p24 (pg/ml), and normalized proviral DNA at week 2 (first measurement in our experiments) without and with the antiviral drug(s) (Fig. 4A). Distribution of model parameters (i.e., $\beta$, $\theta$, $a$, $b$, $p$, $\varepsilon$) along with initial values for variables ($T(0)$, $I(0)$, $L(0)$, $V(0)$) were inferred directly by MCMC computations. The estimated parameters and initial values are listed in tables S4 and S5. Technical details of MCMC computations are summarized below.

Statistical analysis

Package FME (*12*) in R Statistical Software (*13*) was used to infer posterior predictive parameter distributions. The delayed rejection and Metropolis method (*14*) was used as a default computation scheme for FME to perform MCMC computations. MCMC computations for parameter inference were implemented using the pre-defined function modMCMC() in package FME. Convergence of Markov chains to a stationary distribution was required to ensure parameter sets were sampled from a posterior distribution. Only the last 90000 of 100000 chains were used as burn-in. The convergence of the last 90000 chains was manually checked with figures produced by package coda (*15*), a collection of diagnostic tools for MCMC computation. The 95% credible interval shown as a shadowed region in each panel of Fig. 4A was produced from 100 randomly chosen inferred parameter sets and corresponding model predictions. We employed a bootstrap *t*-test (*16*) to quantitatively characterize differences in derived quantities without and with antiviral drug (Fig. 4B). In total, 100000 parameter sets were sampled with replacement from the posterior predictive distributions to calculate the bootstrap t-statistics. To avoid potential sampling bias, the bootstrap *t*-test was performed 100 times repeatedly. The averages of the computed *p*-values were used as indicators of differences.
